# 17α-estradiol Alleviates High-Fat Diet-Induced Inflammatory and Metabolic Dysfunction in Skeletal Muscle of Male and Female Mice

**DOI:** 10.1101/2023.05.30.542870

**Authors:** Matthew P. Bubak, Shivani N. Mann, Agnieszka K. Borowik, Atul Pranay, Albert Batushansky, Samim Ali Mondal, Stephen M. Diodge, Arik Davidyan, Marcelina M. Szczygiel, Fredrick R. Peelor, Sandra Rigsby, Matle Broomfield, Charles I. Lacy, Heather C. Rice, Michael B. Stout, Benjamin F. Miller

## Abstract

Skeletal muscle has a central role in maintaining metabolic homeostasis. 17α-estradiol (17α-E2), a naturally-occurring non-feminizing diastereomer of 17β-estradiol that demonstrates efficacy for improving metabolic outcomes in male, but not female, mice. Despite several lines of evidence showing that 17α-E2 treatment improves metabolic parameters in middle-aged obese and old male mice through effects in brain, liver, and white adipose tissue little is known about how 17α-E2 alters skeletal muscle metabolism, and what role this may play in mitigating metabolic declines. Therefore, this study aimed to determine if 17α-E2 treatment improves metabolic outcomes in skeletal muscle from obese male and female mice following chronic high fat diet (HFD) administration. We hypothesized that male, but not female, mice, would benefit from 17α-E2 treatment during HFD. To test this hypothesis, we used a multi-omics approach to determine changes in lipotoxic lipid intermediates, metabolites, and proteins related to metabolic homeostasis. In male mice, we show that 17α-E2 alleviates HFD-induced metabolic detriments of skeletal muscle by reducing the accumulation of diacylglycerol (DAGs) and ceramides, inflammatory cytokine levels, and reduced the abundance of most of the proteins related to lipolysis and beta-oxidation. In contrast to males, 17α-E2 treatment in female mice had little effect on the DAGs and ceramides content, muscle inflammatory cytokine levels, or changes to the relative abundance of proteins involved in beta-oxidation. These data support to the growing evidence that 17α-E2 treatment could be beneficial for overall metabolic health in male mammals.

## Introduction

Metabolic syndrome is a cluster of conditions that includes obesity, dyslipidemia, hypertension, and insulin resistance.^1, 2^ Skeletal muscle has a central role in maintaining metabolic homeostasis as it accounts for approximately 70-85% of insulin-stimulated glucose disposal^3, 4^ and is the primary tissue responsible for oxidative metabolism. Increases in ectopic lipid accumulation in skeletal muscle can lead to metabolic inflexibility,^5^ which is the reduced capacity to transition between substrates and the accumulation of metabolic byproducts such as diacylglycerol (DAG) and ceramides.^6^ DAG and ceramides are lipotoxic metabolites that interfere with insulin signaling transduction in metabolically active tissues including skeletal muscle^7, 8^ and negatively impact whole-body metabolic homeostasis.^9^ Strategies such as caloric restriction and exercise modulate nutrient-sensing pathways to prevent and/or reverse age- and obesity-related metabolic dysfunction. ^10–13^ However, these strategies are often difficult to maintain due to existing comorbidities.^14, 15^

17α-estradiol (17α-E2) is a naturally-occurring non-feminizing diastereomer of 17β-estradiol (17β-E2) that improves metabolic outcomes in rodents. 17α-E2 supplementation extends median lifespan in male mice when administered during middle^16, 17^ or advanced ages,^18^ which is likely due to metabolic improvements. We and others have shown that in obese and aged male mice, 17α-E2 supplementation reduces caloric intake, ectopic lipid accumulation, and regional adiposity, which was associated with greater glucose tolerance and insulin sensitivity.^19–23^ Importantly, 17α-E2-mediated declines in caloric intake are not necessary for improvements in metabolic parameters,^20^ which suggests that 17α-E2 modulates metabolism through multiple mechanisms such as mTORC2 signaling,^24^ hepatic urea cycling,^25^ and inflammation.^26, 27^ Female mice with intact ovarian function do not respond to 17α-E2.^24–30^ However, female mice with metabolic dysfunction^31^ or that undergo ovariectomy surgeries do gain metabolic benefits from 17α-E2 supplementation.^32^ Even so, chronic administration of 17α-E2 does not protect ovariectomized female mice against age-related metabolic dysfunction.^24–27, 30^ Despite several lines of evidence showing that 17α-E2 treatment improves metabolic parameters in middle-aged obese and old male mice through effects in brain and liver, little is known about how 17α-E2 alters skeletal muscle metabolism.

We have shown that chronic 17α-E2 treatment increases skeletal muscle insulin sensitivity,^21^ but it remains unclear if this occurred through direct actions in skeletal muscle or as a secondary response to improvements in other organ systems. To our knowledge, there is only one report that shows 17α-E2 treatment attenuates age-related muscle loss in male, but not female mice.^30^ Garratt and colleagues found that 17α-E2 treatment in males starting in mid-life increased skeletal muscle amino acid abundance and attenuated skeletal muscle functional declines without altering mTORC1 signaling.^30^ Aged-matched females displayed no benefits to muscle with 17α-E2 treatment in this study.^30^ We have also found that 17α-E2 does not affect skeletal muscle cellular proliferation and proteostatic maintenance when administered late in life,^19^ which further supports the idea that 17α-E2 may elicit benefits in muscle through mTOR-independent mechanisms. It remains unknown if 17α-E2 treatment can improve skeletal muscle metabolomic and lipidomic profiles, and proteostatic maintenance after chronic high-fat diet (HFD) and/or other metabolic challenges.

Therefore, this study aimed to determine if 17α-E2 treatment improves metabolic and proteomic outcomes in skeletal muscle from obese male and female mice following chronic HFD administration. We hypothesized that male, but not female, mice, would benefit from 17α-E2 treatment during a HFD with changes in the mitochondrial proteome to support lipid oxidation and subsequent reductions in DAG and ceramide content. Unexpectedly, we found that 17α-E2 had marked but different beneficial effects within each sex that point to differing mechanisms of action of 17α-E2 between male and female rodents.

## Methods

### Ethical Approval and Study Design

All animal procedures were reviewed and approved by the Institutional Animal Care and Use Committee of the Oklahoma City VA Health Care System. The animals in the current report were part of a larger study published elsewhere.^31^ Body mass and fat mass were presented in this prior report, but we reproduce the data because it provides important context for all of our subsequent downstream analyses. Experimental C57BL/6 mice were bred in-house. At weaning, experimental mice were group-housed by sex and fed standard chow (TestDiet 58YP; 66.6% CHO, 20.4% PRO, 13.0% FAT) until 12 weeks of age. At 12 weeks of age all mice, excluding the chow-fed controls, were fed high-fat diet (HFD; TestDiet 58V8, 35.5% CHO, 18.3% PRO, 45.7% FAT) for 39 weeks (9 months) to induce obesity and metabolic perturbations prior to study initiation. The age-matched chow-fed control mice were evaluated in parallel as a healthy-weight reference group. Two weeks prior to study initiation, all mice were individually housed with ISO cotton pad bedding, cardboard enrichment tubes, and nestlets at 22 ± 0.5°C on a 12:12-hour light-dark cycle. Unless otherwise noted, all mice had ad libitum access to food and water throughout the experimental timeframe. At 12 months of age, mice fed a HFD were randomized into HFD or HFD with the addition of 17α-E2 at 14.4 ppm (HFD+17α-E2, Steraloids, Newport, RI) treatment groups for a 10-week intervention. Body mass and composition (EchoMRI, Houston, TX) were evaluated weekly measurements. Mice were labeled with deuterium oxide (D_2_O, Sigma Aldrich, St. Louis, MO) 10 days prior to sacrifice by injecting the mice with an IP bolus of filtered 99% D_2_O at 30mL/kg and then provided 8% D_2_O in drinking water for 10 days.^19, 34^ At 14-weeks post treatment, mice were fasted for 5-6 hours and euthanized with isoflurane and cardiac puncture. Blood from cardiac puncture was collected into EDTA-lined tubes on ice, and plasma was frozen. Lower limb skeletal muscles were removed, weighed, flash frozen, and stored at −80°C. One gastrocnemius was fixed in 4% paraformaldehyde, cryo-embedded, and stored at −80°C.

### Determination of cellular signaling

For cellular signaling, we used 20–30 mg of homogenized quadriceps muscle for differential centrifugation to save the cytosolic protein fraction. Protein concentration was determined by Peirce 660 assay (cat#: 22660, ThermoFischer Scientific, Walktham, MA) as described previously^35^. Western fractions were processed by the ProteinSimple (San Jose, CA, USA) WES system to separate and visualize proteins according to a standard instrument protocol. The following primary antibodies used were: p-Akt (1:25, Ser473, #4060), Akt (1:50, #4685), α-S6 (1:50, #2217), p-α-S6 (1:25, #4856), AMPK (1:50, #2603), p-AMPK (1:25, #2603), p-4E-BP1 (1:25, Thr37/46, #9459), and 4E-BP1 (1:50, #9452, Cell Signaling, Danvers, MA). Secondary antibodies were included in a WES Master Kit (DM-001, ProteinSimple).

### Skeletal muscle lipid assessments

We used three approaches to assess skeletal muscle lipid species. We first assessed total muscle lipids with Oil Red O. For this assay, one gastrocnemius was embedded in OCT (ThermoFisher, Walham, MA) and cryosectioned into 10 μm sections and slides were stored at −80°C. The Oil Red O (ORO) staining was done according to previously published procedures^36^. In short, the muscle sections were stained with a working solution made using a 1.5:1 ratio of ORO (Sigma Aldrich, St. Louis, MO) to 99% isopropyl alcohol for 10 minutes and then rinsed with distilled water for 30 minutes. Images were taken on a Zeiss 710 confocal microscope (Oberkochen, Germany). Brightfield images were tiled and stitched at 20x for an image of the whole section. ImageJ software was used to quantify the intensity of the stain based on a threshold of 0-172.

Second, we assessed total muscle triacylglycerol content. For this, we homogenized 100mg of quadriceps in 10X (v/w) Cell Signaling Lysis Buffer (Cell Signaling, Danvers, MA) with phosphatase and protease inhibitors (Boston BioProducts, Boston, MA). Total lipid was extracted from this homogenate using the Folch method with a 2:1 chloroform-methanol mixture as commonly described.^37^ A nitrogen drier at room temperature was used to dry the lipid prior to reconstitution in 100 μL of 3:1:1 tert-butyl alcohol-methanol-Triton X-100 solution. Triacylglycerol concentrations were determined using a spectrophotometric assay with a 4:1 Free Glycerol Agent / Triacylglycerol Agent solution (Sigma Triglyceride and Free-Glycerol reagents, St. Louis, MO) as previously described.^38^

Finally, we assessed individual lipid species using a lipidomic approach. For this, we used the NORC Lipidomics Core Laboratory at the University of Colorado-Denver. Approximately 25 mgs of powdered gastrocnemius were lyophilize overnight and we record lyophilized weight for each sample. 150 µL of MilliQ water was added to the lyophilized sample and homogenized for 4’ using bead mill (Quiagen Retsch TissueLyser 2 Bead Mill, Hilden, Germany) at 25 Hz. 800 µL of MilliQ water was added and protein quantification of the homogenate was determined by Peirce 660 assay (cat#: 22660, ThermoFischer Scientific, Walktham, MA). Skeletal muscle homogenate was run according to pervious methods on HPLC-MS/MS^39^ to determine the concentrations of 1,2-diacylglycerols (1-2-DAG), 1,3-diacylglycerols (1-3-DAG), total ceramides (Cer), dihydroceramides (dhCer), 1-O-glucosylceramides (GluCer), 1-O-lactosylceramides (LacCer), and 1-O- galactosylceramides (GlaCer).

### Skeletal muscle inflammation

We used two approaches to determine skeletal muscle inflammation. In the first, total RNA was extracted from 20-30 mg of homogenized tibalis anterior (TA) via Trizol (Life Technologies, Carlsbad, CA) and reverse transcribed to cDNA using the High-Capacity cDNA Reverse Transcription kit (Applied Biosystems, Foster City, CA). Real-time PCR was performed in a QuantStudio 12K Flex Real Time PCR System (Thermofisher Scientific, Waltham, MA) using TaqMan™ Gene Expression Master Mix (Applied Biosystems/Thermofisher Scientific, Waltham, MA) and predesigned gene expression assays with FAM probes from Integrated DNA Technologies (Skokie, Illinois). Target gene expression for IL-6, IL-1α, and TNFα (ThermoFischer, Waltham, MA) was expressed as 2^−ΔΔCT^ by the comparative CT method^40^ and normalized to the expression of HGPRT (ThermoFischer, Waltham, MA) in skeletal muscle.

For our second approach, we assessed concentrations of cytokines in the TA muscle lysate using V-PLEX Mouse Proinflammatory Panel I immunoassays (catalogue no. K15048D-1, Meso Scale Discovery, MD, USA) according to the protocol of the manufacturer. We used 150mg tissue/mL ice cold lysis buffer 500 mM NaCl, 2 mM EDTA, 1% Triton X-100, protease inhibitor cocktail (HALT 100µl/10ml), and 50 mM Tris HCl). Using a probe sonicator set at 40% amplitude we sonicated tissue on ice for 15 seconds, two times. The samples were incubated on a rocker for 30 minutes at 4°C. Following the incubation, we spun the samples for 10 minutes at 20,000g. The concentrations of IFN-γ, IL-10, IL-12p70, IL-1β, IL-2, IL-4, IL-5, IL-6, KC/GRO, and TNF-α were measured in the supernatant and normalized to the protein concentration.

### Assessment of skeletal muscle metabolites

We performed GC-MS-based semi-targeted metabolic profiling for glycolysis, amino acids, and TCA cycle metabolites using an Agilent 7890B gas chromatograph coupled to a 5977A mass spectrometer. Approximately 25 mg of powdered gastrocnemius were homogenized with pre-chilled metal beads using a TissueLyser II (Qiagen, Hilden, Germany) for 40 seconds at 25 Hz. 500 µL of pre-chilled MS-grade methanol was added to the homogenized sample followed by sonication (VWR B1500A-DTH 1.9 L Ultrasonic Cleaner, WR International, LLC, Orange, CA) for 5 minutes. Adonitol (0.2 mg/mL in milli Q water) was added to each sample as internal standard. 250 µL chloroform and 250 µL of milli Q water were added and incubated at 4°C on thermo-shaker (1200 rpm) for 10 minutes. The samples were centrifugated for 10 minutes at max speed (16000 rpm). The supernatant was collected and dried on a SpeedVac (Eppendorf). Dried samples were first derivatized with freshly prepared methoxyaminehydrochloride (115405, MP biomedical, LLC, Irvine, CA) (20 mg/mL in Pyridine, 270407, Sigma-Aldrich, St. Louis MO) on thermo-shaker at 37°C for 90 minutes. Second derivatization was with 35 µL N,O-Bis(trimethylsilyl)trifluoroacetamide (15238- 10x.1ml, Sigma-Aldrich, St. Louis MO) on thermo-shaker at 37°C for 40 minutes. After derivatization, the samples were transferred to glass capped vials and analyzed on GC-MS system with EI source in a scan mode along with the alkane mix (C10 to C24) and an analytical standard mix prepared similarly to samples. Alkane mix (C10 to C24) was used as a retention time standard ladder. The analytical standard mix prepared from standards of pyruvate, lactate, succinic acid, fumaric acid, malic acid, aspartic acid, glutamic acid, citric acid, glucose, fructose-6-phosphate, and glucose-6-phosphate was used for confirmation of these metabolites in the samples. We processed the chromatograms using Mass Hunter Quantitative data analysis software with integrated mass-spectrometry library from the National Institute of Standards and Technology (NIST, Gaithersburg). The metabolites were identified based on external analytical standards (i), and the match of MS spectra and retention indices of the detected targets in the samples with NIST mass-spectrometry library (ii). Samples were randomized and analyzed in two batches and the metabolites relative abundance was calculated by peak area and followed by normalization with sample weight and TIC median. The data from two batches were combined for analysis using integrated exploratory approach to remove the batch effect using ‘limma’ package for R.^41^

### Assessments of protein turnover and content

We performed assessments of bulk protein synthesis and analyzed the change in abundance of individual skeletal muscle proteins. For bulk protein synthesis, approximately 20 mg of quadriceps skeletal muscle was fractionated to measure protein synthesis rates of myofibrillar and mitochondrial proteins according to our previously published procedures.^42–46^ Following tissue fractioning, analytes were prepared for analysis on an Agilent 7890A gas chromatograph coupled to a 5975C mass spectrometer.^42–46^

For individual protein relative abundance, we used 20 µg of gastrocnemius that was lysed in isolation buffer containing 0.02 mM ATP, 100 mM KCl, 10 mM Tris HCl, 10 mM Tris Base, 1 mM MgCl_2_, 0.1 mM EDTA, and 1.5% w/v BSA (fatty acid free). The protein content of the collected supernatant was measured using Pierce 660 Protein Assay (22660, Thermo Fisher Scientific). Protein samples were prepared as previously described using 80 µg of total protein that were spiked with 8 pmol BSA in 1% SDS as an internal standard ^47^. The total protein was precipitated and desalted in 1 mL of acetone overnight at −20°C. The subsequent protein pellet was solubilized in 50 µL Laemmli sample buffer and 20 µg protein was run in a 12.5% SDS-Page gel (BioRad Criterion system). The gels were fixed and stained with Coomassie blue (GelCode blue, Pierce Chemical Company). Each sample was cut from the gel as the entire lane and divided into smaller pieces. A standard in-gel digestion method was used.^48^ Coomassie blue was removed form from the gel and then the gel was reduced in 10 mg/mL DTT, alkylated in 35 mg/mL iodoacetamide, and digested overnight with 1 µg trypsin per sample in 200 µL 10 mM ammonium bicarb. The mixture of peptides was extracted from the gel, evaporated to dryness in a SpeedVac, and reconstituted in 150 μL 1% acetic acid (v/v) for LC-tandem MS analysis. The samples were analyzed using high resolution accurate mass (HRAM) measurements on a ThermoScientific Q-Exactive Plus hybrid quadrupole-orbitrap mass spectrometer using a m/z resolution of 140,000 configured with a splitless capillary column HPLC system (ThermoScientific Ultimate 3000). For discovery proteomics, Mascot Daemon (version 2.8.0) from Matrix Science was used for Mascot database search of tandem mass spectra to identify peptides/proteins.

### Statistical Analyses

Our goal was to test the impact of 17α-E2 within both male and female mice, but not to compare male to female mice. Therefore, within each sex we used a one-way ANOVA (Graphpad Prism 9, San Diego, CA, USA) and R with packages “dplyer”, hmisc”, and “stringer” unless otherwise stated. For changes in body composition over the 12 weeks of treatment (Figure 2A-2F), we used a two-way ANOVA (treatment x time). For the lipid analysis, outliers were removed using R package “Routliers” and mean absolute deviation function.^49, 50^ A Tukey post-hoc analysis was run to determine where differences occurred for the one-way and two-way ANOVAs. For lipid content, statistically significant lipids were considered for further analysis of proteins and their biological relationships using Human Metabolome Data Base (https://hmdb.ca/) and LIPID MAPS Structure Data Bases.^51^ For metabolite abundance the one-way ANOVA was performed on Log10-transformed normalized data under R-environment. For discovery-based proteomics, identified proteins were clustered together within each experimental group. Shared peptides within a group, which we defined as being identified within 50% of the samples, were considered for further analysis of proteins and their biological relationships using Uniprot, Gene Ontology (GO), and PANTHER databases. Further bioinformatic analysis of shared peptides within each group was performed using R programming language (3.4.0) and “dplyer” package.^49^ For the analysis, male and female mice data were subjected to an unsupervised principal component analysis (PCA) separately to evaluate the global effect of 17α-E2 treatment in each sex. PCA was performed on Log10-transformed normalized data using the online tool based on R packages ‘stats’ and ‘heatmap2’.^52^ Results are presented as mean ± standard error (SE) with *p* values less than 0.05 considered to be significant.

**Figure 1:**
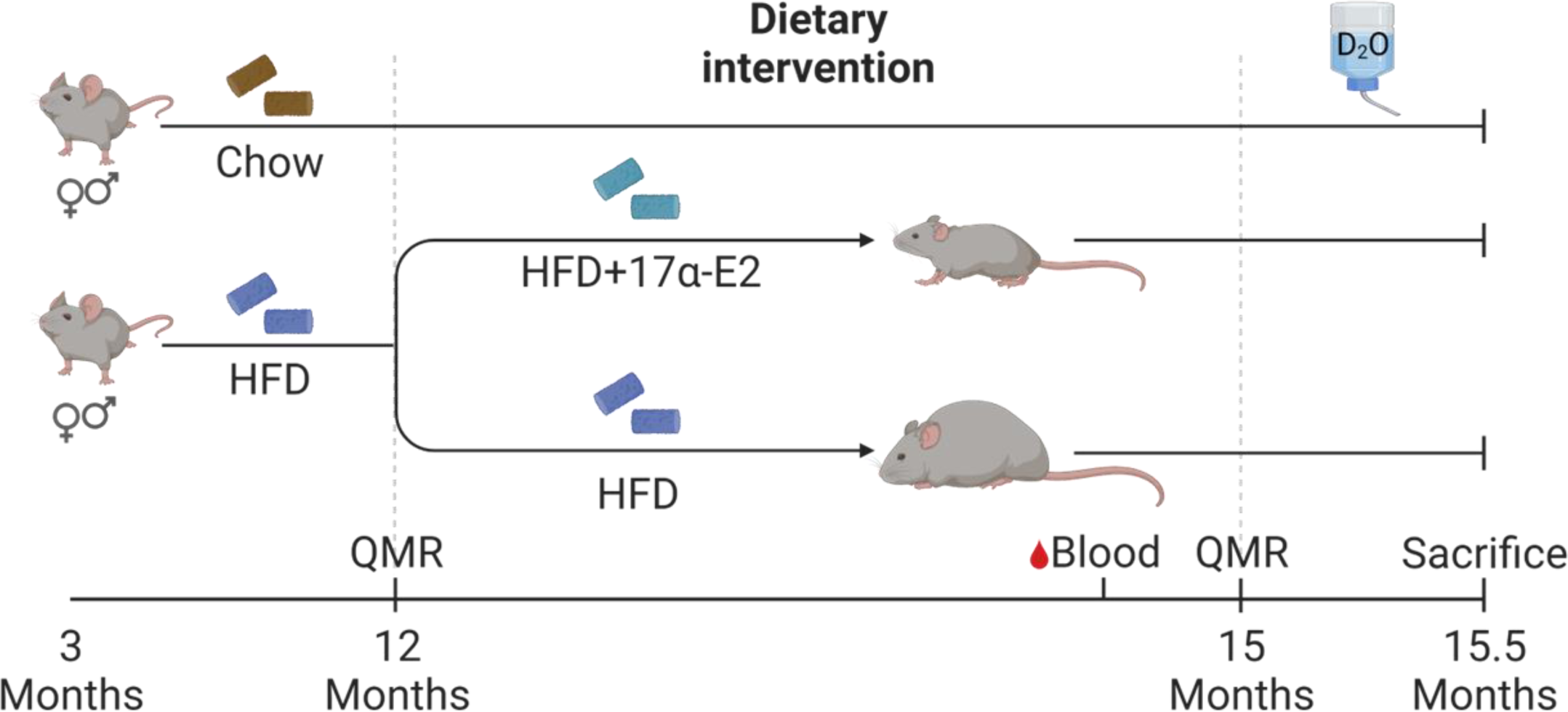
Study design schematic. To understand the effects of 17α-E2 on skeletal muscle in response to a metabolic challenge, male and female mice (3-months) were fed a 45% high-fat diet (HFD). Additionally, male and female (n = 10 and n = 10, respectively) groups of age-matched WT, mice were given a chow diet to act as controls. All mice (age: 12 months) on HFD were randomized into HFD or HFD+17α-E2 (n = 7-10/sex/group) treatment groups for a 3.5-month intervention. All mice underwent deuterium oxide (D_2_O) stable isotope labeling during the last 10 days of the intervention to determine changes to the skeletal muscle myofibrillar and mitochondrial fractions.

**Figure 2:**
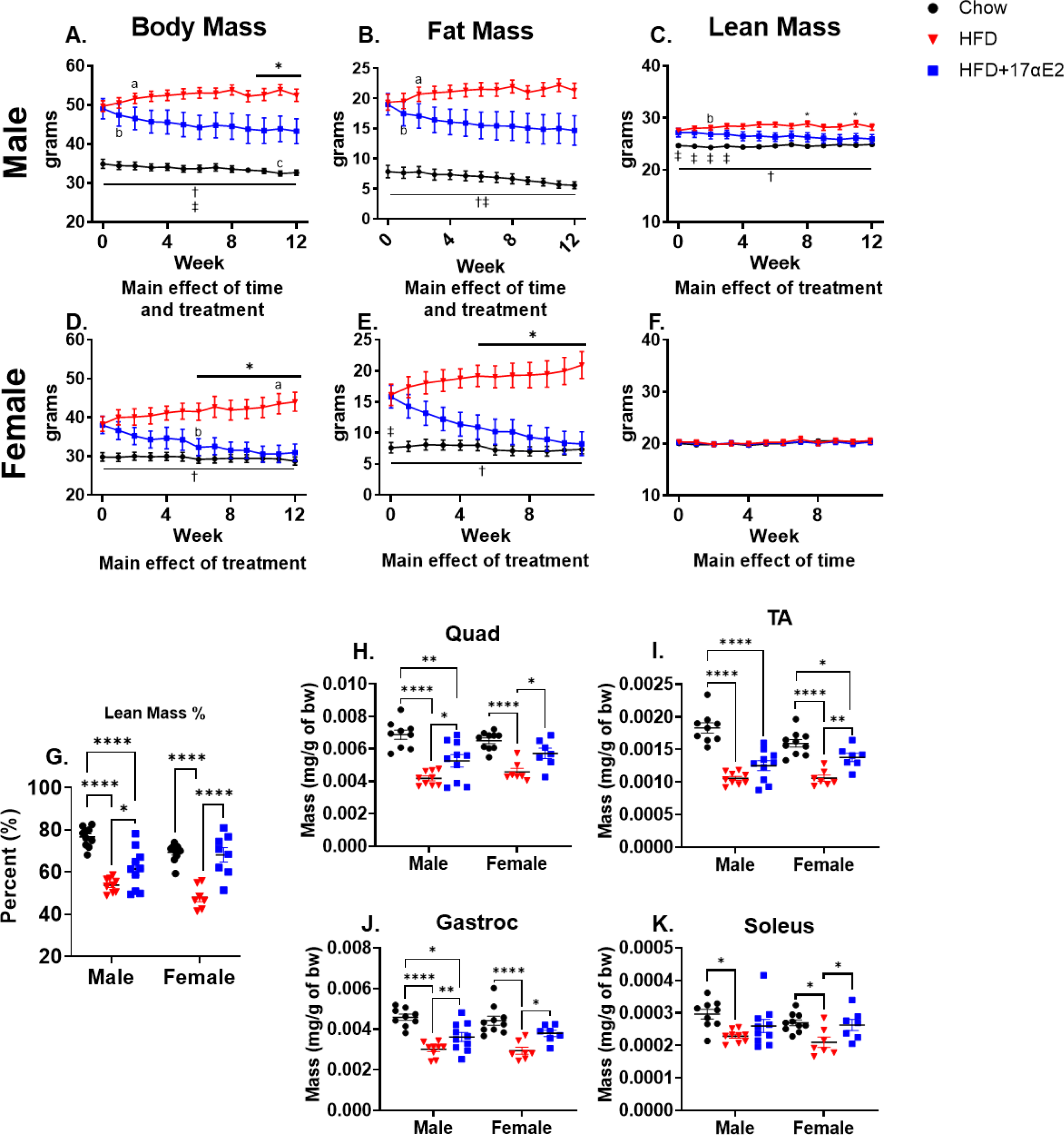
Phenotype description of groups from chow (black), HFD (red), and HFD+17α-E2 (blue). **A-F**) Body composition measures for male (**A-C**) and female mice (**D-F**). Data were analyzed by two-way ANOVA (Treatment x Time) and data are in mean ± SEM from 7–10 mice per group. *p < 0.05 between HFD and HFD+17α-E2, †p < 0.05 between HFD and Chow, ‡p < 0.05 between HDF+17α-E2 and Chow. “a”, “b”, and “c” indicate which timepoint became significantly different (p < 0.05) from week 0 within HFD, HFD+17α-E2, and Chow, respectively. **G**) Lean mass percentage in male and female mice. **H-K**) Skeletal muscle weights normalized to body mass. (**G-K**) Data were analyzed by one-way ANOVA, data are in mean ± SEM from 7–10 mice per group, *p < 0.05, **p < 0.01, ***p < 0.001, and ****p < 0.0001.

## Results

### Body Composition

As shown elsewhere,^31^ body mass and fat mass were found to have a main effect of time and treatment for male mice, and a main effect of treatment in female mice. For fat-free mass, there was a main effect of treatment for male mice, and a main effect of time in the female mice. In male mice, body mass, fat mass and lean mass changed over time (interaction effect, **Figure 2A-C**). In female mice, there was an interaction for body mass and fat mass^31^, but no change in lean mass over time (**Figure 2D-F**). When expressed as percent change, both male and female 17α-E2+HFD had a higher percent lean mass than HFD (**Figure 2G**); in male, but not female mice, 17α-E2+HFD had a lower percent lean mass than Chow (**Figure 2G**). In the male mice, the HFD had lower a relative muscle masses when normalized to body mass in the quadriceps and gastrocnemius compared to the Chow and HFD+17α-E2, while the Chow had greater masses compared to the HFD+17α-E2 (**Figure 2H and 2J**). Additionally, in male mice the tibialis anterior had greater mass in the Chow compared to the HFD and HFD+17α-E2 (**Figure 2I**). Lastly, in male mice the soleus was greater in the Chow compared to the HFD (**Figure 2K**). In female mice the quadriceps, gastrocnemius, and soleus have lower relative masses in the HFD compared to the Chow and HFD+17α-E2 groups (**Figure 2H, J, and K**). Additionally in the female mice, the HFD had lower relative muscle masses when normalized to body mass in the tibialis anterior compared to the Chow and HFD+17α-E2, while the Chow had greater masses compared to the HFD+17α-E2 (**Figure 2I**). However, there was little effect of treatment in both male and female mice when normalized to tibia length (**Supplemental Figure 1**).

### AMPK and mTOR signaling

In the male mice we saw an increase in pAMPK/total levels in the HFD+17α-E2 compared to Chow and HFD, but no difference in female mice (**Figure 3)**. However, female undergoing 17α-E2 treatment had a lower p-AKT/total ratio than the Chow. These data show that 17α-E2 treatment had little effect on downstream targets of mTOR pathway and AMPK signaling in males or females on a HFD (**Figure 3 and Supplemental Figure 2**).

**Figure 3:**
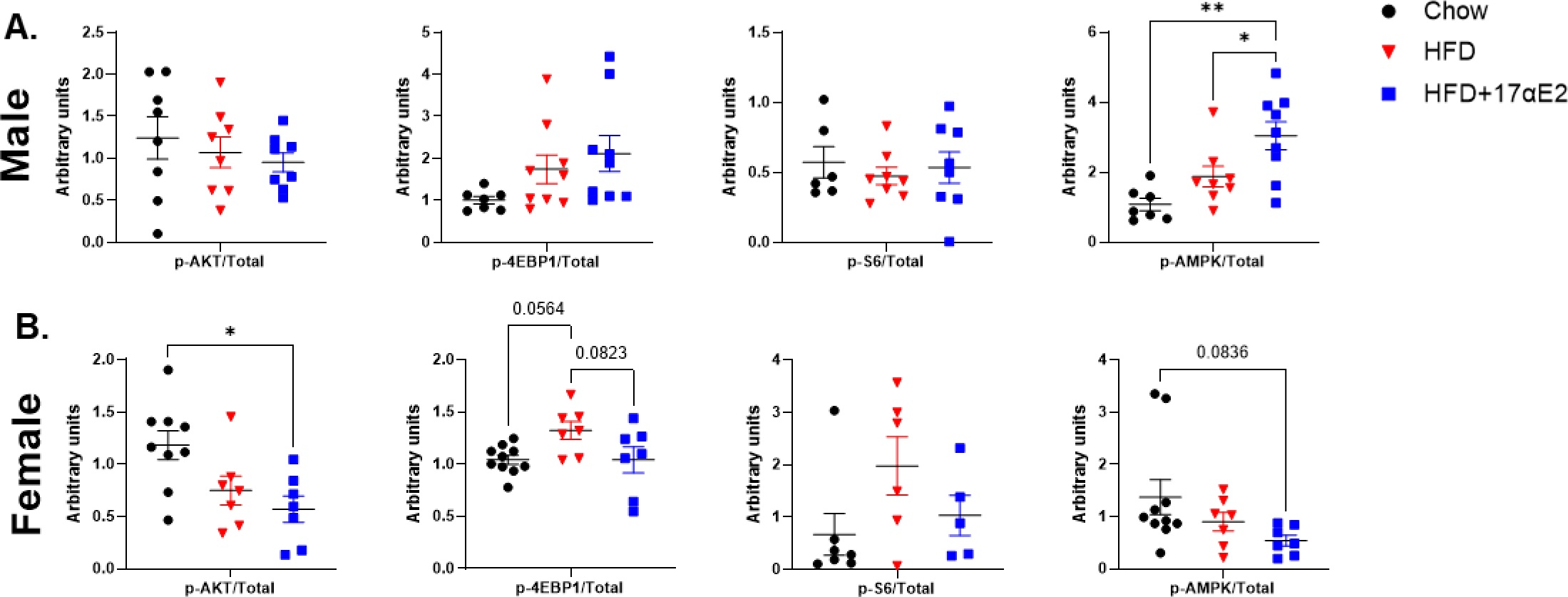
Results for p-AKT:AKT, p-4E-BP1:4E-BP1, p-S6:S6, and p-AMPKα:AMPKα ratios in **A**) male and **B**) female mice, respectively. Representative blots of p-AKT, AKT, p-S6, S6, 4E-BP1, p-4E-BP1, p-AMPKα, and AMPKα from Chow, HFD, and HFD+17α-E2. Data are expressed as mean ± SEM of 6–10 mice per treatment group and were analyzed by one-way ANOVA, *p < 0.05 and **p < 0.01.

### Muscle lipid-related outcomes

Male HFD had greater in Oil Red O staining than Chow and a trend toward greater staining in the HFD+17α-E2 compared to Chow (**Figure 4A and B**). Female HFD+17α-E2 was greater than Chow, with a trend toward higher concentration in HFD versus Chow (**Figure 4A and B**). Compared to Chow, male HFD and HFD+17α-E2 had higher quadriceps triacylglycerol concentration (**Figure 4C**). In female mice, HFD had higher triacylglycerol concentration than both Chow and HFD+17α-E2, while HFD+17α-E2 was not different from Chow (**Figure 4C**).

**Figure 4:**
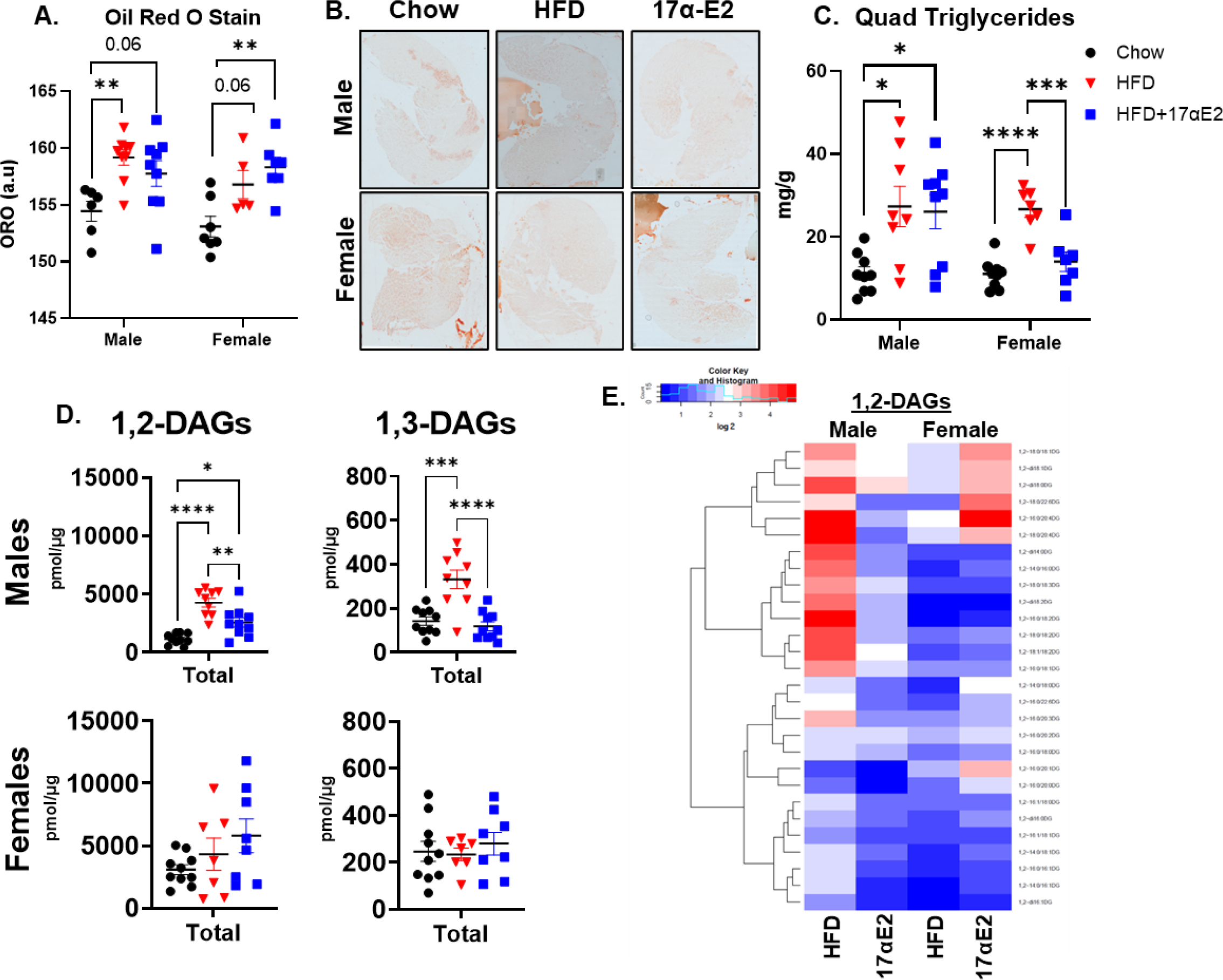
**A**) Concentration of triacyclglycerols in the quadriceps muscle. **B**) Oil-red-O staining intensity in gastrocnemius in arbitrary units (a.u). **C**) Representative images of Oil-Red-O stain. **D**) Total 1,2-DAGs and 1,3-DAG content of the gastrocnemius muscle. **E**) Individual 1,2-DAGs represented as fold change from Chow. All lipids shown are significantly different in male and/or female mice. *p < 0.05, **p < 0.01, ***p < 0.001, and ****p < 0.0001. Values are in mean ± SEM from 7–10 mice per treatment group and were analyzed by one-way ANOVA.

In the gastrocnemius of male mice, there were greater total 1,2-DAGs and 1,3-DAGs in HFD versus Chow, while HFD+17α-E2 was less than HFD for both (**Figure 4D**). In contrast, there were no differences between the treatment groups in female mice across the majority of total saturated, unsaturated, monounsaturated, and polyunsaturated FAs (**Supplemental Figures 3-9**). When presented as a heat map, there are striking differences between male and female mice in how HFD and HFD+17α-E2 change 1,2 DAG profiles (**Figure 4E**). Additionally, there appears to be changes that are specific to male or female mice and the direction of those changes in six lipid species **(Figure 4E)**. The Human Metabolite Data Base was used to determine the physiological function of these six DAGs and revealed that these DAGs are intermediates for plasmalogen synthesis, de novo triacylglycerol synthesis, and beta-oxidation of long chain fatty acids (**supplemental table 1 and 2**). In contrast to the changes in 1,2-DAG and 1,3-DAG concentrations, there were no changes with treatment in total ceramides for male and female mice (**Supplemental Figure 5**). When broken down by subclasses of ceramides, HFD male mice had greater total and unsaturated dhCeramides and LacCeramides compared to Chow and HFD+17α-E2 (**Supplemental Figure 6 and 9**).

### Skeletal muscle inflammation

Our results show that the male HFD mice had greater mRNA expression of IL-6, IL-1α, and TNFα compared to the Chow and HFD+17α-E2, but there were no differences between the Chow and HFD+17α-E2 (**Figure 5A**). In the female mice, HFD did not change mRNA expression of any cytokine (**Figure 5A**). When examining cytokine concentrations in the muscle, male HFD had greater concentration of IL-10 compared to Chow and HFD+17α-E2, while the HFD+17α-E2 was greater compared to Chow. The male HFD had greater concentration of IL-12p70 and KCGRO compared to Chow, less IL-6 concentration compared to Chow, and HFD+17α-E2 had lower TNFα concentrations compared to HFD. In the female mice, the HFD had greater concentration of IL-10 compared to Chow and HFD+17α-E2. Also, the female HFD+17α-E2 had a greater IL-6 concentration compared to Chow. There were no differences among the groups for the cytokines IFNγ, IL-1B, IL-2, IL-4, IL-5 in male or female mice and no differences in TNFα or IL-12p70 in the female mice (**Figure 5B** and **supplemental figure 10**). IFNγ and IL-4 were below the detection range of the stand curve (**supplemental figure 10**).

**Figure 5:**
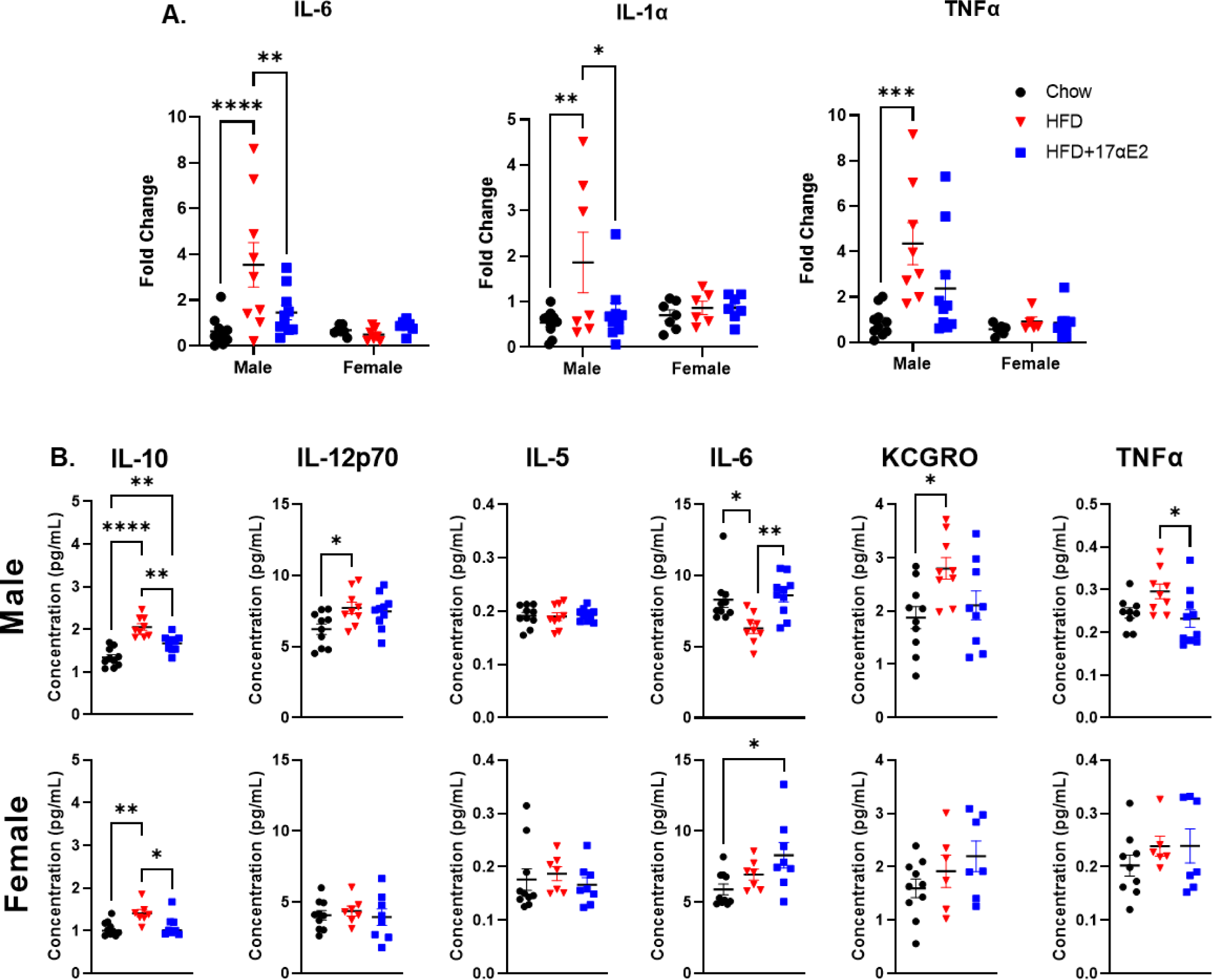
**A**) mRNA expression of inflammatory cytokines in the TA muscle. **B**) The pro-inflammatory cytokine and chemokine protein concentrations in the plantaris muscle. Values are in mean ± SEM from 7–10 mice per treatment group and were analyzed by one-way ANOVA,*p < 0.05, **p < 0.01, and ****p < 0.0001.

### Skeletal muscle metabolites

Across the principal component 1 (PC_1_), there was a separation based on diet between the Chow versus HFD and HFD+17α-E2 (regardless of 17α-E2 treatment) in male mice, but a smaller separation of HFD and HFD+17α-E2. Across PC_2_, there is a small separation between the HFD and HDF+17α-E2, however, only a small percentage of the variance is explained by 17α-E2 treatment (**Figure 6A**). Conversely, female groups were not clearly separated (**Figure 6B**). Our results show that HFD and HFD+17α-E2 had greater alanine, succinic acid, and serine abundance in both male and female mice compared to Chow, while glycine, beta alanine, aminomalonic acid and adenosine were altered in abundance in response to HFD and HFD+17α-E2 in male mice only (**Figure 6C and Supplemental Figure 11**). Tyrosine was elevated in the HFD compared to the HFD+17α-E2 in female mice (**Figure 6D**). The female HFD mice had a greater abundance of taurine and tyrosine compared to Chow (**Figure 6D and Supplemental Figure 11**). However, there were only three metabolites glucose-6-phosphate, threonine, and succinic acid that were different in the HFD compared to the HFD+17α-E2 in male mice (**Figure 6C, Supplemental Figure 11**). Glucose-6-phosphate and threonine were lower in HFD compared to HFD+17α-E2 in male mice while succinic acid was higher in HFD compared to HFD+17α-E2 (**Figure 6C and Supplemental Figure 11**). We did not observe similar change in these metabolites between HFD and HFD+17α-E2 in female mice (**Figure 6D and Supplemental Figure 11**).

**Figure 6:**
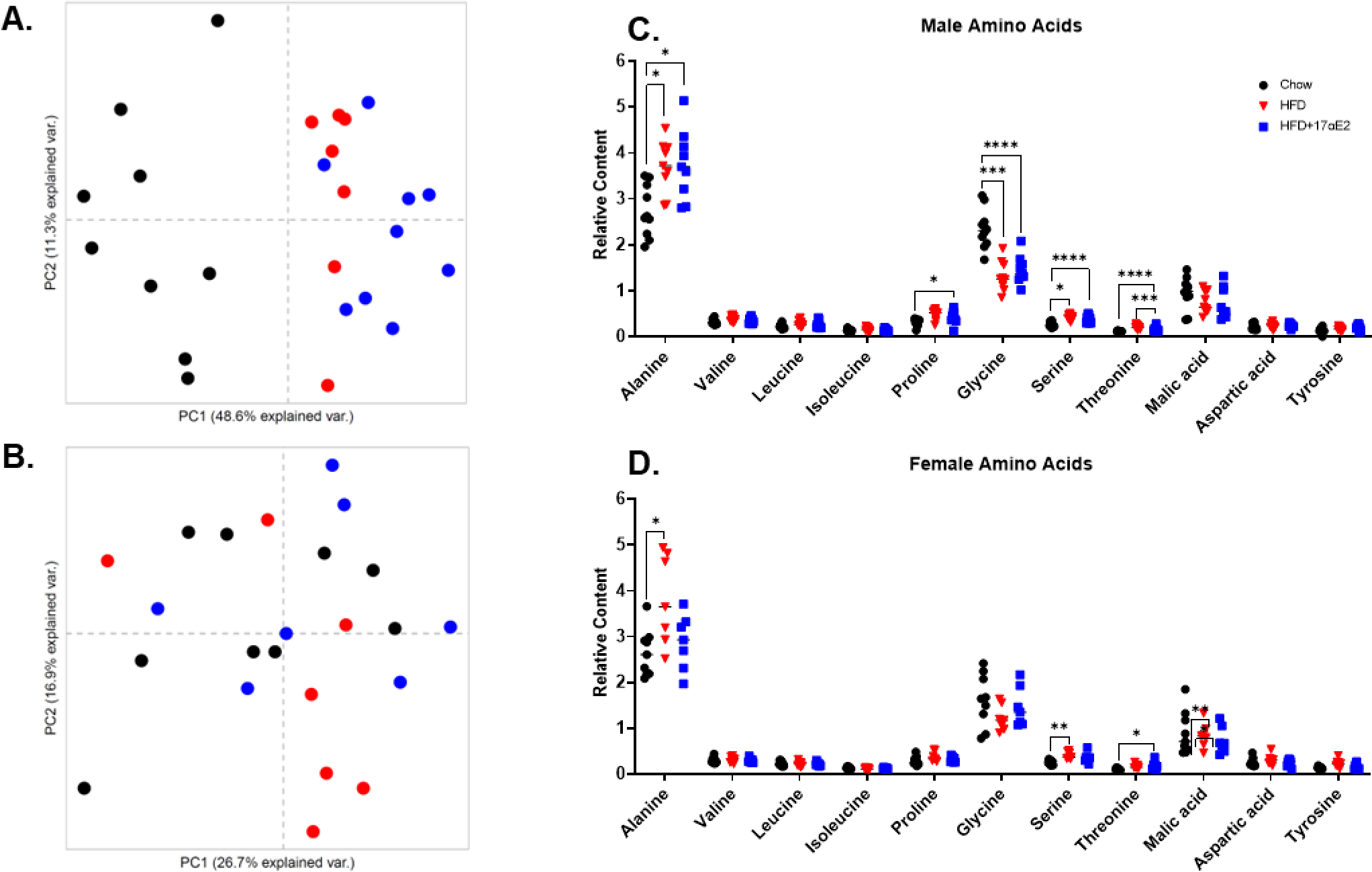
PCA plots for metabolite abundance in **A**) male and **B**) female mice. Relative abundance of amino acid content in gastrocnemius of **C**) male and **D**) female mice. Values are in mean ± SEM from 7–10 mice per treatment group and were analyzed by one-way ANOVA,*p < 0.05, **p < 0.01, ***p < 0.001, and ****p < 0.0001.

### Protein turnover and proteomic analysis

In the quadriceps, the myofibrillar fractional synthesis rate (FSR) was greater in the HFD and HFD+17α-E2 compared to Chow of male mice. There were no differences in the myofibrillar FSR among the female mice (**Figure 7A**). In the mitochondrial fraction, the male mice had greater FSR in the HFD and HFD+17α-E2 compared to Chow. The female HFD mice had greater mitochondrial FSR compared to the Chow and HFD+17α-E2 (**Figure 7B**).

**Figure 7:**
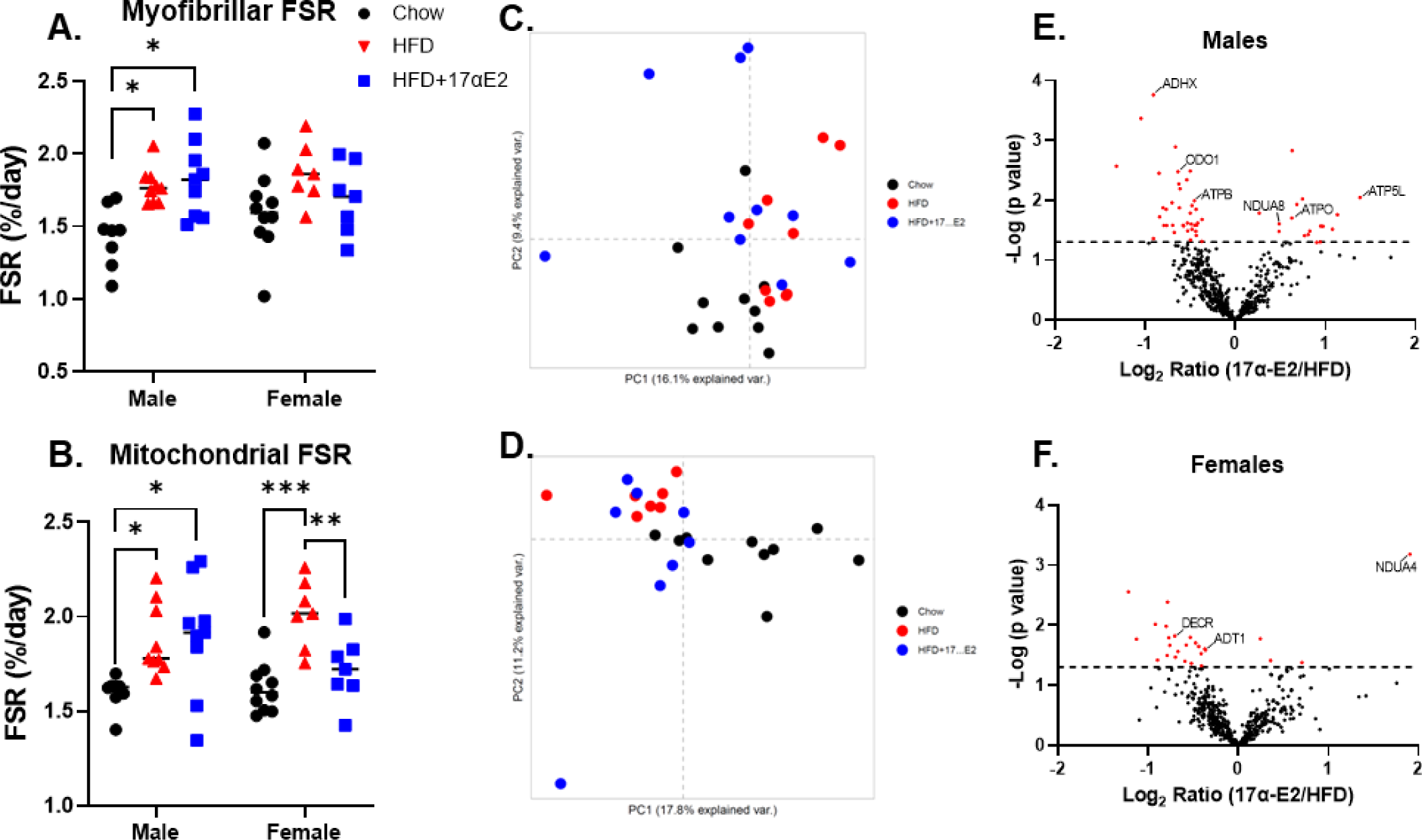
**A**) Myofibrillar and **B**) mitochondrial fraction synthesis rates (%/Day). PCA for **C**) males and **D**) female mice. Volcano plot representing the relative difference in protein abundance between HFD+17α-E2 and HFD of **E**) male and **F**) female mice. These data are from 7–10 mice per treatment group and were analyzed by one-way ANOVA, *p < 0.05, **p < 0.01, and ***p < 0.001.

In the male mice, the PCA plot shows limited clustering among the groups. HDF+17α-E2 partially clusters with the HFD, thus a low amount of variance is explained by the 17α-E2 (**Figure 7C**). In the female mice, the PCA shows no clear separation (**Figure 7D**). In the male mice, the relative abundance of 52 of 618 identified proteins were significantly different (>log2-fold change of 1.3) in the HFD+17α-E2 compared to HFD, while in the female mice 26 of 586 identified proteins were significantly different in the HFD+17α-E2 compared to HFD (**Figure 7E and 7F**). Of the 52 proteins identified as being significantly different between the male HFD+17α-E2 and HFD mice, 20 were involved in mitochondrial and beta oxidation processes, 17 were structural and/or contractile proteins, nine were related to synthetic processes, and seven had other biological functions. In the female mice, 26 proteins identified as being significantly different between the male HFD+17α-E2 and HFD mice, eight were involved in mitochondrial and beta oxidation processes, three were structural and/or contractile proteins, five were related to synthetic processes, and 10 had other biological functions. The abundance and primary functions of the significantly different individual proteins are listed in **Supplemental table 3.** The HFD+17α-E2 had greater abundance of mitochondrial proteins involved in the mitochondrial membrane ATP synthase and the oxidization of long chain fatty acids, carbohydrates and amino acids (ADHX, ODO1, and ATPB) compared to HDF. Interestingly, there was also a greater abundance of proteins involved in mitochondrial membrane ATP synthase and NADH dehydrogenase (NDUA8 ATOP, and ATP5L) in the HFD+17α-E2 compared to HFD (**Figure 7E**). These significantly different proteins were group by GO terms the enriched pathways involved ATP metabolic process (GO:0046034), oxidative phosphorylation (GO:0006119), proton motive force-driven mitochondrial ATP synthesis (GO:0042776), and carbohydrate catabolic process (GO:0016052, **Supplemental table 4**). In female HFD+17α-E2 mice there was a reduced abundance of mitochondrial-related proteins (6 of 7) such as those involved in beta-oxidation and ATP synthesis (DECR and ADT1) and a great abundance of NDUA4, a component of the cytochrome c oxidase which drives oxidative phosphorylation, in the HFD+17α-E2 compared to HFD (**Figure 7F**). These proteins are involved in energy production and beta oxidations thus it appears that 17α-E2 treatment alters mitochondrial beta oxidation and skeletal muscle metabolism. When we ran GO on the significantly different proteins in the female group, there was no enrichment in biological pathways.

## Discussion

We tested the hypothesis that 17α-E2 would be protective against HFD-induced metabolic derangements in skeletal muscle of male, but not female, mice. As reported elsewhere,^31^ 17α-E2 improves body composition in both sexes, but also improves metabolic parameters in the skeletal muscle of male mice on a HFD. We show that 17α-E2 alleviates HFD-induced metabolic detriments of skeletal muscle by reducing the accumulation of bioactive lipids, inflammatory cytokine levels, and reduced the abundance of most of the proteins related to lipolysis and beta-oxidation. In contrast to males, 17α-E2 treatment in female mice had little effect on the muscle inflammatory status, the majority of bioactive lipid intermediates, or changes to the relative abundance of proteins involved in beta-oxidation. These data add further support to the growing evidence that 17α-E2 treatment could be beneficial for overall metabolic health in male mammals.

The expansion of lipid storage in adipose tissue leads to the release of pro-inflammatory cytokines^53^ and ectopic lipid storage in other tissues such as skeletal muscle.^54^ This storage of lipids in muscle can lead to impairment of insulin-stimulated glucose uptake,^55^ contributing to overall insulin resistance. We explored whether the beneficial effects of 17α-E2 treatment on insulin sensitivity in male mice were partly due to improvements in skeletal muscle. To ensure an obese phenotype in the mice, we fed a HFD for approximately 9 months prior to treatment with 17α-E2. As was our goal, the male mice had approximately double the fat mass compared to chow when we started 17α-E2 treatment. Although lean mass was slightly increased in the HFD at the start of treatment, the percent lean mass was greater at the end of treatment. Therefore, there was less lean mass, and potentially muscle mass, to help maintain metabolic flexibility. After 12 weeks of 17α-E2 treatment, there were improvements in percent lean mass despite continued HFD, supporting previous studies in male mice.^21, 23^ Although there were minimal differences in absolute muscle masses (**supplemental figure 1**) at the end of the treatment period, the muscle mass normalized to body mass was significantly improved with 17α-E2 treatment (**Figure 1H-K**), thus improving the potential to maintain metabolic flexibility.

The HFD also increased ectopic lipid deposition in male mice as evidenced by higher Oil Red O staining and muscle triglycerides. Under low flux conditions, as with sedentary behavior, incomplete oxidation of triacylglycerol via lipases results in the accumulation of bioactive lipid metabolites such as DAGs and ceramides.^7^ These lipid intermediates are thought to mediate of lipid-induced insulin resistance.^8, 9, 56^ Consistent with this idea, the higher levels of triglycerides in male mice were matched by higher overall levels of DAGs, but not ceramides. Our lipidomic analyses showed that both 1,2 and 1,3-DAGs were higher with HFD compared to chow. In addition, the DAG 18:0, which has been linked to insulin resistance,^57^ was elevated with HFD in male mice. In the male mice, these changes in lipid species were associated with significantly higher expression of the pro-inflammatory cytokines IL-6, IL-1α, and TNFα, which are linked to whole-body insulin sensitivity.^58^ Interestingly, IL-10, which is an anti-inflammatory cytokine, was elevated in both sexes on the HFD which is likely due to compensatory mechanisms. Treatment with 17α-E2 during HFD had several interesting changes to lipids in skeletal muscle. The most striking was that even though total lipids and TG remained higher than Chow, there was a significant decrease in 1,2 and 1,3 DAGs, including 18:0, compared to HFD. These differences were matched by significantly lower expression of IL-6 and IL-1α. These findings support that 17α-E2 in male mice has positive changes in lipid profiles in skeletal muscle, potentially improve insulin sensitivity in muscle.

Our findings in female mice were partially unexpected. First, treatment with 17α-E2 completely reversed the HFD-induced increase in fat mass so that there were no longer differences compared to Chow. Like male mice, 17α-E2 treatment resulted in higher percentages of lean mass and muscle mass normalized to body mass. With HFD, female mice also had significantly greater muscle triglycerides and a trend (p=0.06) toward higher total lipids. Unlike male mice, HFD in female mice did not increase 1,2 or 1,3 DAGs or expression of any pro-inflammatory cytokines. In addition, the muscle concentration of cytokines, which would likely include contributions from the systemic environment, also showed minimal changes with HFD in female mice. It is thought that compared to male adipose tissue, female adipose tissue has a greater capacity to expand and store lipids thus minimizing ectopic lipid stored.^59^ Although our data showed increased lipids in female muscle, it is tempting to speculate that high turnover was maintained in the lipid pools, thus minimizing the accumulation of DAGs. HFD+17α-E2 had lower muscle triglyceride levels compared to HFD, but there were no minimal changes to DAGs or expression of cytokines in muscle because they were not greater with HFD to begin with. Therefore, our data showed some overall benefits on body composition of female mice that were fed a prolonged HFD, but there were minimal improvements in muscle because the muscle was remarkably resistant to HFD-induced changes to begin with.

There is one more interesting finding with the regard DAGs that is worth noting. There were six 1,2 DAG species (18:0/18, 18:1, 18:0, 18:0/22, 16:0/20, and 18:0/20) that had essentially opposite responses to HFD and 17α-E2 treatment. These species were higher in the male HFD compared to Chow but were not different from Chow with HFD+17α-E2. In female HFD, these species were not different between Chow and HFD, but were greater with HFD+17α-E2. While in male mice these changes mirror those of other 1,2 DAGs, in female mice they are the only ones that increased with 17α-E2 treatment. According to the Human Metabolome Data Base, these six 1,2-DAGs are involved in de novo triacylglycerol biosynthesis, plasmalogen synthesis, and mitochondrial beta-oxidation of long-chain saturated fatty acids (**supplemental tables 1 and 2**).

We wanted to determine if changes in lipid deposition and inflammation impacted muscle metabolism. To do so, we used a semi-targeted metabolomic profiling approach that focused on 45 metabolites. From our PCA, we saw that male mice are clearly separated by Chow versus HFD, while female mice had no separation. Interestingly, 17α-E2 treatment did not result in any further separation in male mice. We focused on amino acid concentrations because a previous study showed that in aged male mice, 17α-E2 had greater skeletal muscle amino acid abundance compared to controls while aged female mice had a lower abundance of amino acids compared to controls.^30^ Although we found some changes in amino acids concentrations in both sexes, the changes were not unidirectional or consistent with HFD and HFD+17α-E2, limiting our ability to make broad conclusions about the impacts on amino acid metabolism.

In male mice, HFD+17α-E2 had increased AMPK activation compared to both Chow and HFD. A previous study in male mice without HFD did not see an increase in AMPK activation of skeletal muscle, although there was an increase in white adipose tissue.^21^ Activation of AMPK signaling is thought to be the primary mechanism of the improvements in insulin sensitivity by the anti-diabetic drug metformin.^60^ In the case of metformin, this results in a lowering of hepatic glucose output and improvements in glucose uptake in muscle.^61–63^ Although inhibition of mTOR often follows activation of AMPK, we did not find any changes in mTOR signaling in the muscle of male or female mice. This finding further supports the notion that the positive effects of 17α-E2 are not through the inhibition of mTOR like other healthspan-extending treatments.^21, 24, 30^

Previously, we examined the impact of 17α-E2 on protein turnover in older mice.^19^ In that study, there was a lack of changes in protein turnover, although a group that was pair-fed to be deficient in calories equal to the reduced intake from 17α-E2 treatment did have changes in protein turnover. In the current study, male mice had an increase in myofibrillar protein synthesis with HFD as we have demonstrated in other studies previously.^64, 65^ Surprisingly, there was not an increase in mTOR signaling that is often present in skeletal muscle with obesity. It is important to note that the measurements of protein synthesis were made after many months of Chow or HFD and in the last 2 weeks of a 12-week 17α-E2 treatment. Therefore, at this point, lean mass had stabilized, and the synthesis rates likely indicative a steady state and not acute remodeling. If protein synthesis is higher, and lean mass and muscle masses are maintained, there must also be a higher rate of protein breakdown. This higher turnover rate would indicate improved maintenance of proteostasis in the HFD+17α-E2. There was also an increase in mitochondrial protein synthesis with HFD in male mice. We have previously shown increases in mitochondrial protein synthesis during HFD,^64, 66, 67^ but these studies are shorter in duration and were thought to be compensatory changes for the diet. Interestingly, rates of mitochondrial protein synthesis were also higher in the HFD+17α-E2 group. Our proteomic analysis showed extensive differences in the mitochondrial proteome between HFD and HFD+17α-E2 group. Therefore, although both HFD and HFD+17α-E2 had greater mitochondrial protein FSR than Chow, these rates are resulting in different proteins; likely compensatory to HFD in one case, and corrective in the other. This speculation is supported by a lower relative abundance of the majority of individual mitochondrial proteins involved in beta-oxidation and energy production in HFD+17α-E2 compared to HFD (**Figure 8B, supplemental table 3 and 4**); many of which are downstream of ERα signaling.^68–71^

In female mice, it is interesting that HFD did not increase myofibrillar protein synthesis. As far as we are aware, this is the first study demonstrating this difference between the sexes in myofibrillar protein synthesis rates to HFD. This finding also adds to the overall picture that skeletal muscle of female mice is protected from HFD-induced detriments. As opposed to myofibrillar protein synthesis, the mitochondrial protein synthesis was higher than chow with HFD. When female mice had 17α-E2 during HFD, the mitochondrial protein synthesis was not different from Chow. Since our proteomics analysis also showed a dearth of changes in mitochondria-related protein abundance between HFD and HFD±17α-E2, there was likely no substantive differences with treatment, perhaps because none were needed. With treatment, there was a rapid reduction in body and fat mass of female mice and no differences in triacylglycerol concentrations, DAG or ceramide content, or inflammatory cytokines between the Chow and 17α-E2 treatments. Therefore, we speculate that the muscle of female mice is resistant to detrimental effects of HFD, thus not deriving muscle-specific benefits from 17α-E2 treatment.

There are a couple of limitations to our study. First, we could not assess early changes to treatment with 17α-E2 since we focused on outcomes 12 weeks into treatment. Although we were interested in long-term changes, there are early changes from 17α-E2 treatment that would have been interesting to assess from a remodeling perspective. Second, we did not assess the impact of 17α-E2 concurrent with the onset of HFD. In that sense, we examined the therapeutic potential of 17α-E2 rather than the potential to prevent HFD-induced changes in metabolism. That said, it is generally thought that reversing conditions is more challenging than prevention, thus demonstrating the great benefits of 17α-E2 treatment. Finally, we were limited in our assessments of metabolic flux. Adding other tracers needed for flux measurements would be challenging analytically since the D_2_O that was planned for turnover measurements. However, true flux measurements would allow us to further understand the positive changes in muscle lipid species.

## Conclusions

Our study adds to the growing literature that the healthspan benefits of 17α-E2 are likely mediated through positive metabolic changes. In the present study we focus on skeletal muscle and show that the positive effects differ between sexes. In male mice, there were direct benefits on the skeletal muscle that resulted in positive changes in lipid species, decreased inflammation, and likely improved remodeling to maintain proteostasis. In female mice, the impact of HFD on skeletal muscle was minimal to begin with, thus 17α-E2 treatment had fewer benefits in the muscle. However, the 17α-E2 treatment greatly reduced fat mass and likely has other systemic benefits as a result. Therefore, contrary to our hypothesis, 17α-E2 benefit female mice, although these may be minimal directly on skeletal muscle. Future studies will focus on better understanding changes in metabolic flux in skeletal muscle with 17α-E2 treatment.

## Acknowledgements

Some images were created with BioRender.com. We would like to acknowledge the Center for Biomedical Data Sciences at the Oklahoma Medical Research Foundation for the assistance with bioinformatics and figure generation.

## Data Availability

The data underlying this article will be shared on reasonable request to the corresponding author.

---

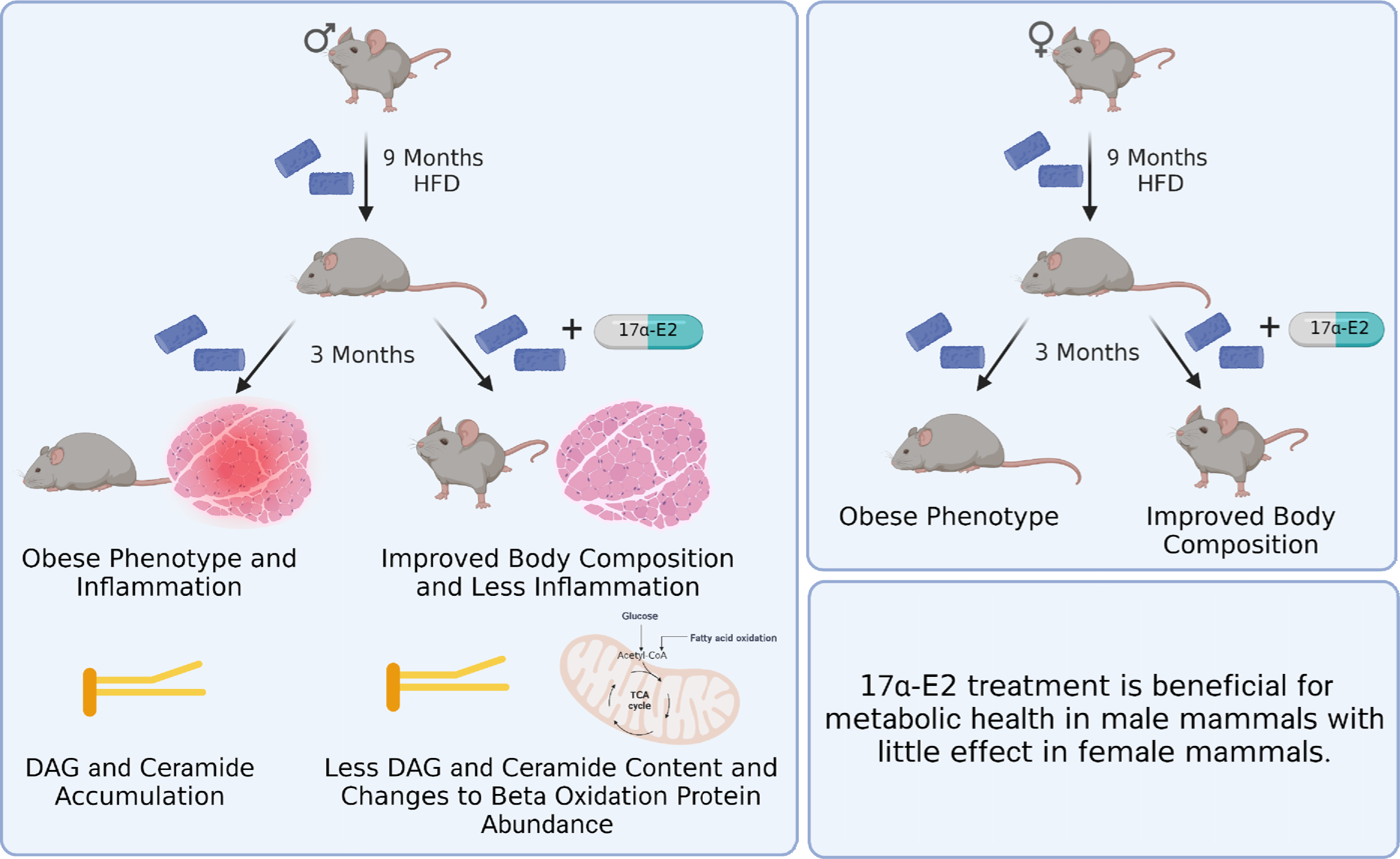

