## Supplemental Figure 1 for "17α-estradiol Alleviates High-Fat Diet-Induced Inflammatory and Metabolic Dysfunction in Skeletal Muscle of Male and Female Mice"

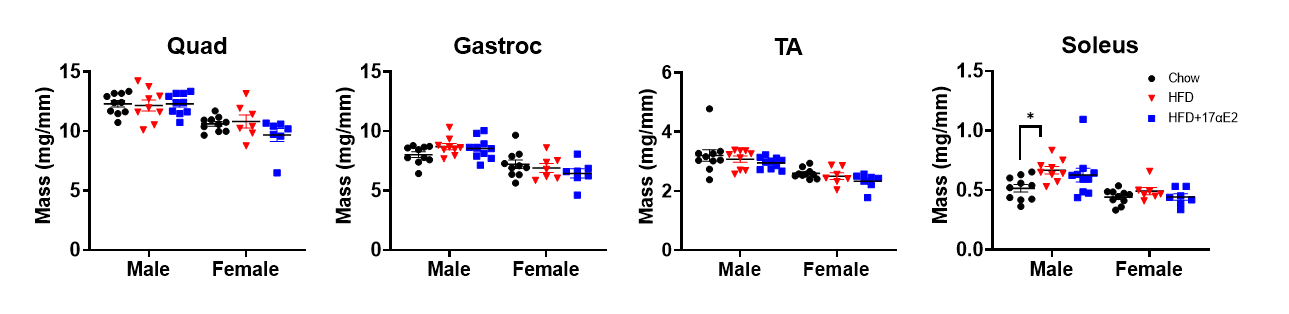

**Supplemental Figure 1:** Skeletal muscle masses normalized to tibia length. Data were analyzed by one-way ANOVA, data are in mean ± SEM from 7–10 mice per group,*p < 0.05.
