## Supplemental Figure 2 for "17α-estradiol Alleviates High-Fat Diet-Induced Inflammatory and Metabolic Dysfunction in Skeletal Muscle of Male and Female Mice"

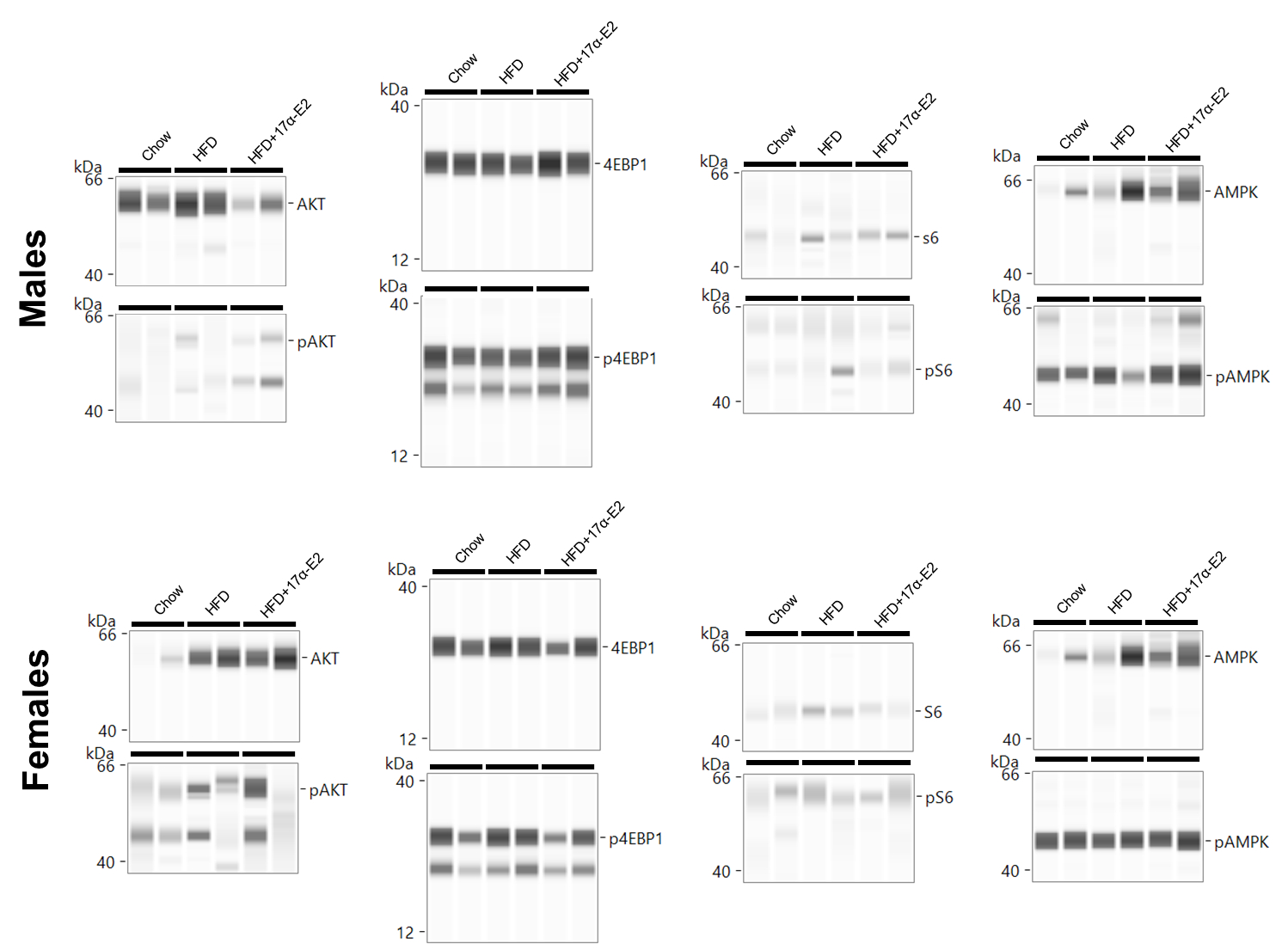

**Supplementary Figure 2:** Representative blots of p-AKT, AKT, p-S6, S6, 4E-BP1, p-4E-BP1, p-AMPKα, and AMPKα from Chow, HFD, and HFD+17α-E2.
