## Supplemental Figure 3 for "17α-estradiol Alleviates High-Fat Diet-Induced Inflammatory and Metabolic Dysfunction in Skeletal Muscle of Male and Female Mice"

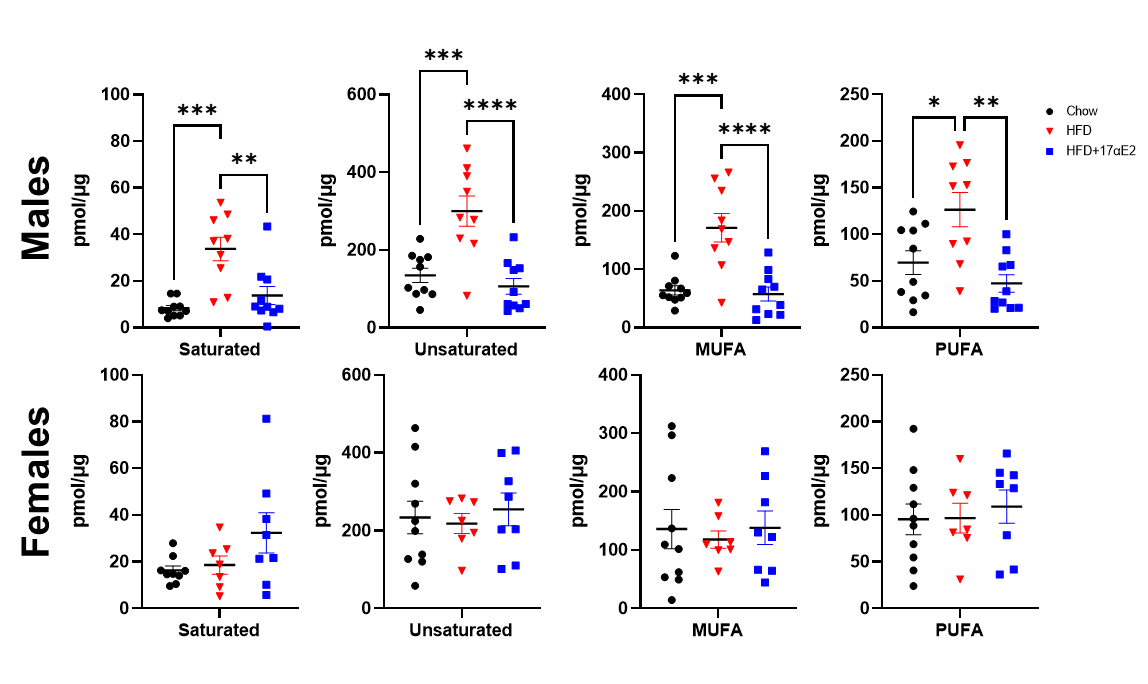

**Supplemental Figure 3:** 1,2-DAGs for males and females represented as saturated, unsaturated, monounsaturated, and polyunsaturated (PUFA) fatty acids. *p < 0.05, **p < 0.01, ***p < 0.001, and ****p < 0.0001. Values are in mean ± SEM from 7–10 mice per treatment group and were analyzed by one-way ANOVA.
