## Supplemental Figure 5 for "17α-estradiol Alleviates High-Fat Diet-Induced Inflammatory and Metabolic Dysfunction in Skeletal Muscle of Male and Female Mice"

**Supplemental Figure 5:** Ceramide content for males and females represented as total, saturated, unsaturated, monounsaturated, and polyunsaturated (PUFA) fatty acids. *p < 0.05 and **p < 0.01. Values are in mean ± SEM from 7–10 mice per treatment group and were analyzed by one-way ANOVA.

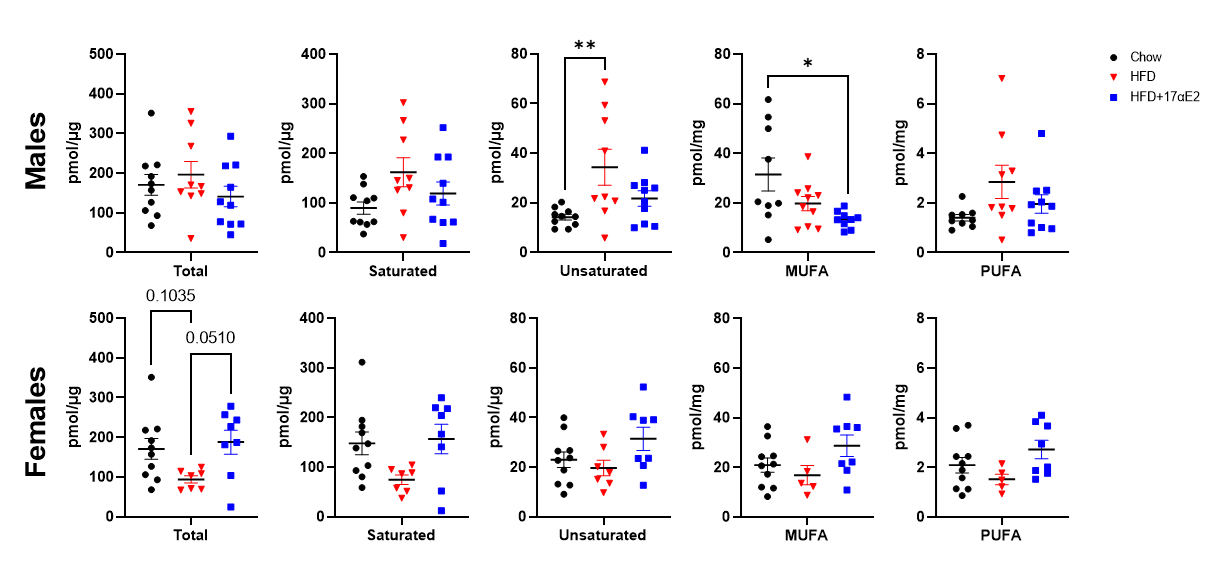
