## Supplemental Figure 6 for "17α-estradiol Alleviates High-Fat Diet-Induced Inflammatory and Metabolic Dysfunction in Skeletal Muscle of Male and Female Mice"

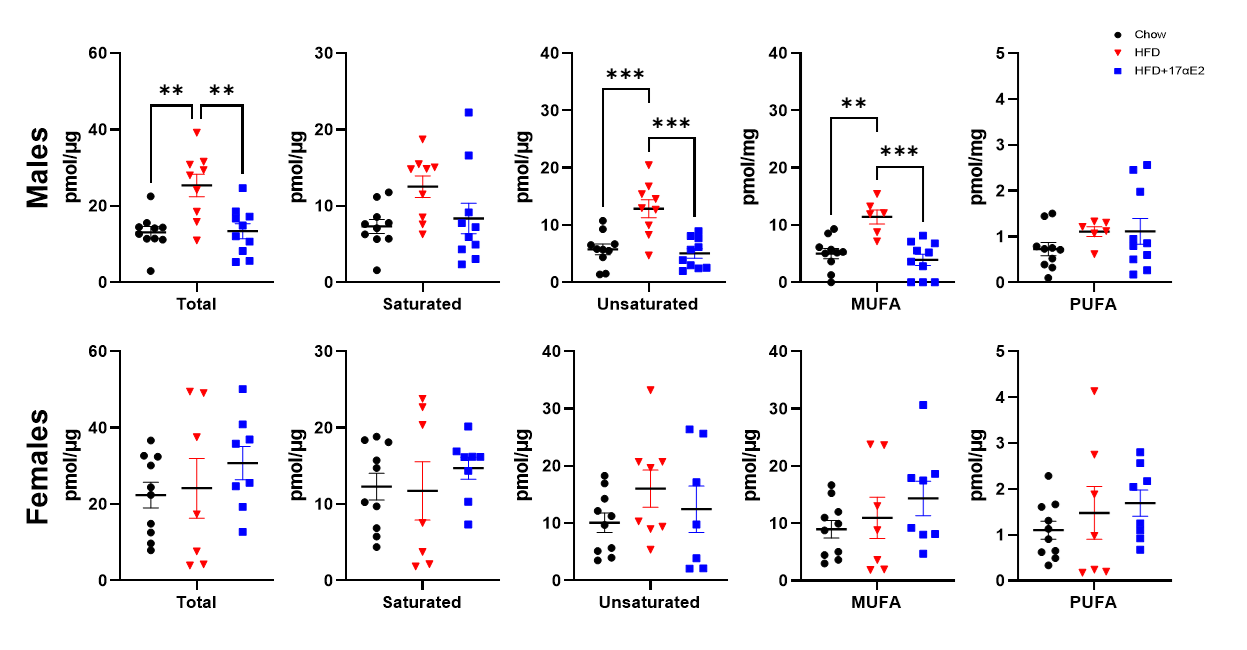

**Supplemental Figure 6:** Dihydroceramide content for males and females represented as total, saturated, unsaturated, monounsaturated, and polyunsaturated (PUFA) fatty acids. **p < 0.01 and ***p < 0.001. Values are in mean ± SEM from 7–10 mice per treatment group and were analyzed by one-way ANOVA.
