## Supplemental Figure 8 for "17α-estradiol Alleviates High-Fat Diet-Induced Inflammatory and Metabolic Dysfunction in Skeletal Muscle of Male and Female Mice"

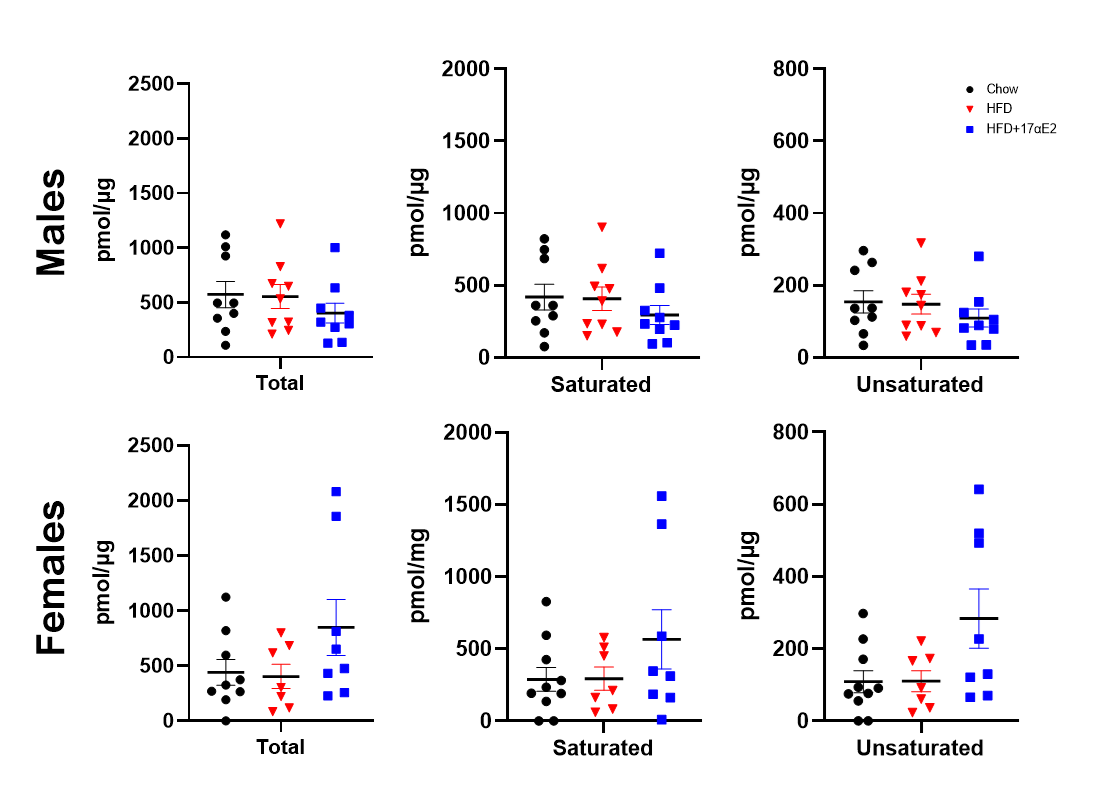

**Supplemental Figure 8:** Glucosylceramides content for males and females represented as total, saturated, unsaturated, monounsaturated, and polyunsaturated (PUFA) fatty acids. Values are in mean ± SEM from 7–10 mice per treatment group and were analyzed by one-way ANOVA.
