## Supplemental Figure 10 for "17α-estradiol Alleviates High-Fat Diet-Induced Inflammatory and Metabolic Dysfunction in Skeletal Muscle of Male and Female Mice"

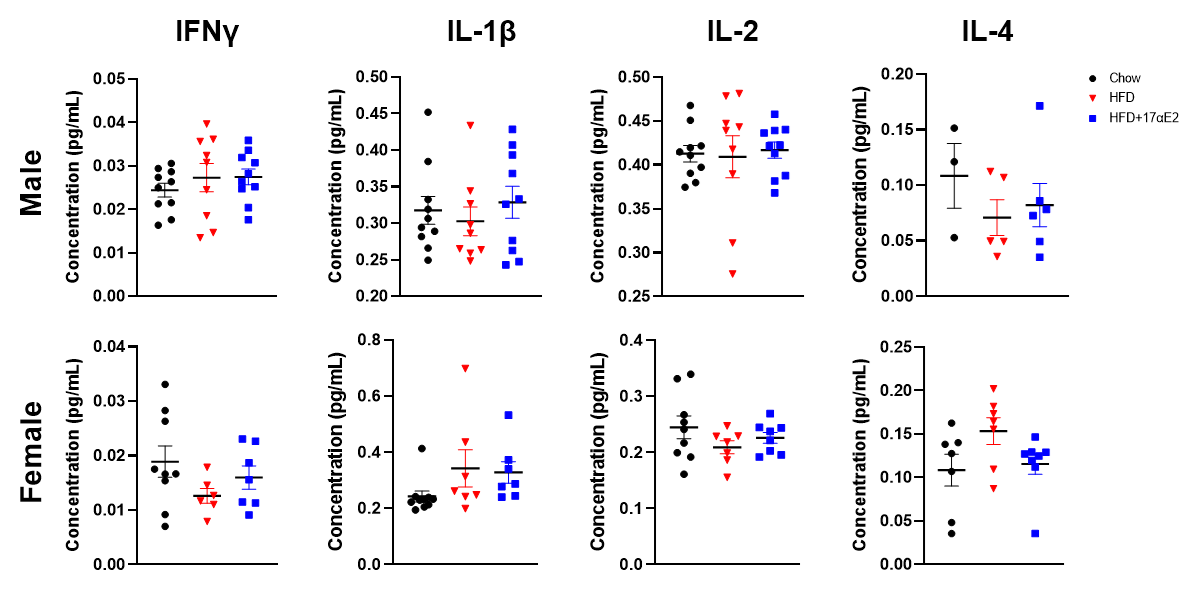

**Supplemental Figure 10:** The pro-inflammatory cytokine and chemokine concentrations in the plantaris muscle. IL-4 samples were blow detectable ranges. Values are in mean ± SEM from 7–10 mice per treatment group and were analyzed by one-way ANOVA.
