## Supplemental Figure 11 for "17α-estradiol Alleviates High-Fat Diet-Induced Inflammatory and Metabolic Dysfunction in Skeletal Muscle of Male and Female Mice"

**Supplemental Figure 11:** Fold change relative to chow & transformed to log2 scale from 7–10 mice per treatment group and were analyzed by one-way ANOVA.

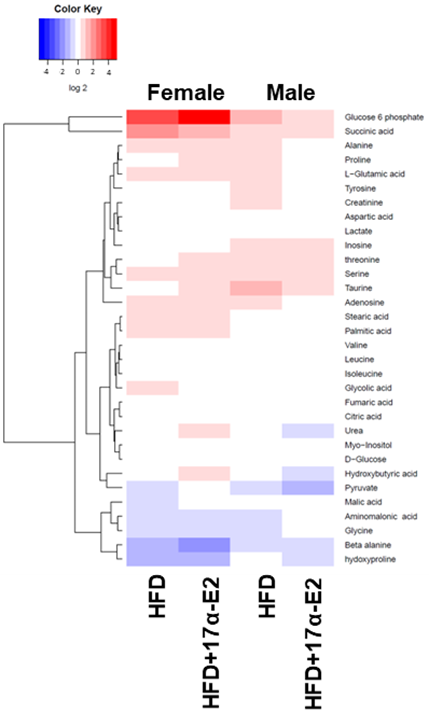
