## Supplemental Table 1 for "17α-estradiol Alleviates High-Fat Diet-Induced Inflammatory and Metabolic Dysfunction in Skeletal Muscle of Male and Female Mice"

**Supplemental table 1**: Individual 1,2-DAGs represented as fold change from control group from 7–10 mice per treatment group and were analyzed by one-way ANOVA.

| **Dag Species** | **Human Metabolome ID** | **Biological Processes** | **Female** | |  | **Male** | |
| --- | --- | --- | --- | --- | --- | --- | --- |
|  |  |  | **HFD** | **17α-E2** |  | **HFD** | **17α-E2** |
| 1,2-di18:0/18:1DG | HMDB07160 | De Novo Triacylglycerol Biosynthesis | 1.268 | 1.781 |  | 1.802 | 1.378 |
| 1,2-di18:1 | HMDB00207 | NA | 1.178 | 1.635 |  | 1.511 | 1.326 |
| 1,2-di18:0 | HMDB0000827 | Mitochondrial Beta-Oxidation of Long Chain Saturated Fatty Acids | 1.237 | 1.617 |  | 2.031 | 1.569 |
| 1,2-di18:0/22:6DG | HMDB0007179 | De Novo Triacylglycerol Biosynthesis | 0.609 | 1.987 |  | 1.543 | 0.592 |
| 1,2-di16:0/20:4DG | HMDB07118 | Plasmalogen Synthesis | 1.402 | 2.271 |  | 2.2 | 1.092 |
| 1,2-di18:0/20:4DG | HMDB0007176 | De Novo Triacylglycerol Biosynthesis | 1.107 | 1.684 |  | 2.219 | 0.592 |
