## Supplemental Table 2 for "17α-estradiol Alleviates High-Fat Diet-Induced Inflammatory and Metabolic Dysfunction in Skeletal Muscle of Male and Female Mice"

**Supplemental table 2**: 1,2-DAG content (pmol/mg) of dry tissue of the six individual DAGs that had inverse response to 17α-E2 treatment in male and female mice. Data are in mean ± standard deviation.

|  |  | **1,2-DAG Content pmol/mg of dry tissue** | | | | | |
| --- | --- | --- | --- | --- | --- | --- | --- |
|  |  | **18:0/18:1DG** | **di18:1DG** | **di18:0DG** | **18:0/22:6DG** | **16:0/20:4DG** | **18:0/20:4DG** |
| **Male** | **HFD** | 225.7 ± 56.8 | 1068.0 ± 331.1 | 10.2 ± 3.9 | 48.3 ± 20.1 | 20.5 ± 7.5 | 365.5 ± 217 |
|  | **HFD+17αE2** | 168.2 ± 119.1 | 939.7 ± 692.6 | 7.4 ± 6.0 | 25.0 ± 15.5 | 9.5 ± 4.1 | 118.3 ± 34.6 |
|  | **Chow** | 64.8 ± 43.6 | 374.7 ± 274.0 | 2.5 ± 1.7 | 16.6 ± 9.2 | 4.5 ± 3.5 | 78.5 ± 54.5 |
| **Female** | **HFD** | 296.7 ± 254.1 | 1433.1 ± 1236.8 | 13.5 ± 11.2 | 32.7 ± 36.7 | 17.6 ± 17.25 | 320.8 ± 327.2 |
|  | **HFD+17αE2** | 423.3 ± 366.8 | 1968.3 ± 1761.5 | 17.5 ± 14.8 | 85.1 ± 79.6 | 32.1 ± 21.9 | 478.4 ± 434.5 |
|  | **Chow** | 123.2 ± 67.3 | 633.6 ± 337.0 | 5.7 ± 4.1 | 21.5 ± 8.8 | 6.7 ± 2.8 | 148.9 ± 139.7 |
