## Supplemental Table 3 for "17α-estradiol Alleviates High-Fat Diet-Induced Inflammatory and Metabolic Dysfunction in Skeletal Muscle of Male and Female Mice"

**Supplemental table 3**: The relative abundance of the significantly different individual proteins is listed in supplementary table 4. Data are represented as fold change from 7–10 mice per treatment group and were analyzed by one-way ANOVA.

| **Male Mice** | | | |
| --- | --- | --- | --- |
| **Protein** | **Uniprot ID** | **Log2 Fold Change 17α-E2/HFD** | **-log (p value)** |
| ADHX_MOUSE | P28474 | -0.91036 | 3.761954 |
| PRPS1_MOUSE | Q9D7G0 | -1.04679 | 3.367543 |
| JPH1_MOUSE | Q9ET80 | -0.66474 | 2.890421 |
| MYL3_MOUSE | P09542 | 0.632066 | 2.828566 |
| TLN1_MOUSE | P26039 | -1.32193 | 2.568958 |
| KCC2A_MOUSE | P11798 | -0.50148 | 2.48825 |
| ODO1_MOUSE | Q60597 | -0.63627 | 2.477556 |
| TYW4_MOUSE | Q8BYR1 | -0.848 | 2.451611 |
| MYH3_MOUSE | P13541 | -0.53959 | 2.342371 |
| DHDH_MOUSE | Q9DBB8 | -0.62795 | 2.271971 |
| TRFE_MOUSE | Q921I1 | -0.61372 | 2.192059 |
| ATP5L_MOUSE | Q9CPQ8 | 1.386498 | 2.044312 |
| MGDP1_MOUSE | Q9D967 | 0.745305 | 2.020953 |
| ATPB_MOUSE | P56480 | -0.45523 | 1.991059 |
| DPYL2_MOUSE | O08553 | -0.70302 | 1.957149 |
| TNNC2_MOUSE | P20801 | 0.683248 | 1.930813 |
| ACOT2_MOUSE | Q9QYR9 | -0.48067 | 1.910625 |
| ARK72_MOUSE | Q8CG76 | -0.80239 | 1.875333 |
| MYH1_MOUSE | Q5SX40 | -0.62016 | 1.874909 |
| MYH4_MOUSE | Q5SX39 | -0.76785 | 1.849151 |
| KPYM_MOUSE | P52480 | -0.43732 | 1.848937 |
| ATP4A_MOUSE | Q64436 | 0.265894 | 1.78566 |
| MYH8_MOUSE | P13542 | -0.50021 | 1.783649 |
| COF1_MOUSE | P18760 | 1.133856 | 1.757782 |
| EFTU_MOUSE | Q8BFR5 | -0.84199 | 1.720675 |
| ATPO_MOUSE | Q9DB20 | 0.62781 | 1.700362 |
| ENOA_MOUSE | P17182 | -0.36907 | 1.677595 |
| EF1G_MOUSE | Q9D8N0 | -0.41835 | 1.619499 |
| ACTN2_MOUSE | Q9JI91 | -0.53179 | 1.616867 |
| NDUA8_MOUSE | Q9DCJ5 | 0.487728 | 1.604185 |
| UBA1_MOUSE | Q02053 | -0.48962 | 1.601973 |
| EF1A2_MOUSE | P62631 | -0.45193 | 1.583027 |
| PLEC_MOUSE | Q9QXS1 | -0.58108 | 1.582645 |
| MYPC2_MOUSE | Q5XKE0 | -0.43 | 1.582429 |
| TITIN_MOUSE | A2ASS6 | -0.69594 | 1.580127 |
| 2AAA_MOUSE | Q76MZ3 | -0.79474 | 1.578462 |
| VIME_MOUSE | P20152 | -0.76369 | 1.577082 |
| ACTN3_MOUSE | O88990 | -0.5753 | 1.572969 |
| NDUAC_MOUSE | Q7TMF3 | 0.948679 | 1.568556 |
| ATP5I_MOUSE | Q06185 | 0.974618 | 1.566438 |
| ACYP2_MOUSE | P56375 | 1.078003 | 1.514591 |
| AT1A2_MOUSE | Q6PIE5 | -0.48006 | 1.514548 |
| NIPS2_MOUSE | O55126 | -0.52978 | 1.487663 |
| RACK1_MOUSE | P68040 | 0.828234 | 1.48617 |
| GSTM1_MOUSE | P10649 | -0.43949 | 1.480238 |
| TRPM7_MOUSE | Q923J1 | 0.485427 | 1.478352 |
| DECR_MOUSE | Q9CQ62 | -0.67993 | 1.463164 |
| SODC_MOUSE | P08228 | 0.809693 | 1.425483 |
| MYOM1_MOUSE | Q62234 | -0.43419 | 1.412514 |
| NDUA5_MOUSE | Q9CPP6 | 0.769819 | 1.40789 |
| KAPCB_MOUSE | P68181 | -0.91058 | 1.359916 |
| SPEG_MOUSE | Q62407 | -0.49943 | 1.334982 |
| PHB2_MOUSE | O35129 | -0.37292 | 1.310904 |
| **Female Mice** | | | |
| **Protein** | **Uniprot ID** | **Log2 Fold Change 17α-E2/HFD** | **-log (p value)** |
| NDUA4_MOUSE | Q62425 | 1.907815104 | 3.181115 |
| RS16_MOUSE | P14131 | -1.22080617 | 2.556424 |
| MYH1_MOUSE | Q5SX40 | -0.786557895 | 2.38426 |
| K2C72_MOUSE | Q6IME9 | -0.920928429 | 2.014035 |
| KLH41_MOUSE | A2AUC9 | -0.801693197 | 1.981966 |
| DECR_MOUSE | Q9CQ62 | -0.705231523 | 1.82281 |
| MYH8_MOUSE | P13542 | -0.532376431 | 1.796695 |
| TRFE_MOUSE | Q921I1 | -0.772708779 | 1.792635 |
| CASQ1_MOUSE | O09165 | 0.244673869 | 1.773838 |
| AMPL_MOUSE | Q9CPY7 | -1.130969394 | 1.767665 |
| KPB1_MOUSE | P18826 | -0.474495733 | 1.704169 |
| HXK2_MOUSE | O08528 | -0.579990843 | 1.666694 |
| MYH4_MOUSE | Q5SX39 | -0.760559643 | 1.665888 |
| LDHB_MOUSE | P16125 | -0.439853819 | 1.646276 |
| KCC2G_MOUSE | Q923T9 | -0.372732466 | 1.606372 |
| ADT1_MOUSE | P48962 | -0.364249725 | 1.586097 |
| RS27A_MOUSE | P62983 | -0.669577111 | 1.561458 |
| MYH3_MOUSE | P13541 | -0.412530248 | 1.527565 |
| ESTD_MOUSE | Q9R0P3 | -0.79017104 | 1.496087 |
| PCCB_MOUSE | Q99MN9 | -0.699986682 | 1.46773 |
| HIBCH_MOUSE | Q8QZS1 | -0.89566334 | 1.418745 |
| ANXA6_MOUSE | P14824 | 0.357787512 | 1.412457 |
| NU1M_MOUSE | P03888 | -0.591194454 | 1.396931 |
| NACAM_MOUSE | P70670 | 0.707819249 | 1.377589 |
| CBR1_MOUSE | P48758 | -0.521199495 | 1.360334 |
| TITIN_MOUSE | A2ASS6 | -0.406769883 | 1.321245 |
