## Supplemental Table 4 for "17α-estradiol Alleviates High-Fat Diet-Induced Inflammatory and Metabolic Dysfunction in Skeletal Muscle of Male and Female Mice"

**Supplemental table 4**: GO Ontology database for male mice of the significantly different individual proteins listed in supplementary table 2.

| Analysis Type: | PANTHER Overrepresentation Test (Released 20221013) | | | | | | |
| --- | --- | --- | --- | --- | --- | --- | --- |
| Annotation Version and Release Date: | GO Ontology database DOI: 10.5281/zenodo.6799722 Released 2022-07-01 | | | | | | |
| Analyzed List: | upload_1 (Mus musculus) | | | | | | |
| Reference List: | Mus musculus (all genes in database) | | | | | | |
| Test Type: | FISHER |  |  |  |  |  |  |
| Correction: | FDR |  |  |  |  |  |  |
| GO biological process complete | Mus musculus REFLIST (21997) | Number | Expected | Plus/Minus | Fold Enrichment | Raw P Value | FDR |
| skeletal myofibril assembly (GO:0014866) | 8 | 2 | 0.02 | + | > 100 | 2.52E-04 | 4.27E-02 |
| ATP metabolic process (GO:0046034) | 160 | 13 | 0.39 | + | 33.72 | 1.73E-16 | 2.73E-12 |
| purine ribonucleoside triphosphate metabolic process (GO:0009205) | 187 | 13 | 0.45 | + | 28.85 | 1.16E-15 | 9.16E-12 |
| ribonucleoside triphosphate metabolic process (GO:0009199) | 192 | 13 | 0.46 | + | 28.1 | 1.60E-15 | 8.43E-12 |
| purine nucleoside triphosphate metabolic process (GO:0009144) | 193 | 13 | 0.47 | + | 27.96 | 1.71E-15 | 6.74E-12 |
| nucleoside triphosphate metabolic process (GO:0009141) | 210 | 13 | 0.51 | + | 25.69 | 4.80E-15 | 1.51E-11 |
| purine nucleotide metabolic process (GO:0006163) | 371 | 15 | 0.89 | + | 16.78 | 1.16E-14 | 3.04E-11 |
| ribose phosphate metabolic process (GO:0019693) | 387 | 15 | 0.93 | + | 16.09 | 2.10E-14 | 4.72E-11 |
| purine-containing compound metabolic process (GO:0072521) | 404 | 15 | 0.97 | + | 15.41 | 3.84E-14 | 7.56E-11 |
| purine ribonucleotide metabolic process (GO:0009150) | 356 | 14 | 0.86 | + | 16.32 | 1.46E-13 | 2.56E-10 |
| ribonucleotide metabolic process (GO:0009259) | 376 | 14 | 0.91 | + | 15.45 | 2.99E-13 | 4.71E-10 |
| nucleotide metabolic process (GO:0009117) | 477 | 15 | 1.15 | + | 13.05 | 3.93E-13 | 5.63E-10 |
| nucleoside phosphate metabolic process (GO:0006753) | 484 | 15 | 1.17 | + | 12.86 | 4.82E-13 | 6.33E-10 |
| muscle system process (GO:0003012) | 253 | 12 | 0.61 | + | 19.69 | 1.22E-12 | 1.48E-09 |
| generation of precursor metabolites and energy (GO:0006091) | 338 | 13 | 0.81 | + | 15.96 | 1.60E-12 | 1.80E-09 |
| nucleobase-containing small molecule metabolic process (GO:0055086) | 537 | 15 | 1.29 | + | 11.59 | 2.05E-12 | 2.15E-09 |
| muscle contraction (GO:0006936) | 197 | 11 | 0.47 | + | 23.17 | 2.18E-12 | 2.14E-09 |
| phosphate-containing compound metabolic process (GO:0006796) | 1772 | 22 | 4.27 | + | 5.15 | 3.69E-11 | 3.42E-08 |
| carbohydrate derivative metabolic process (GO:1901135) | 931 | 17 | 2.24 | + | 7.58 | 3.81E-11 | 3.34E-08 |
| ATP biosynthetic process (GO:0006754) | 83 | 8 | 0.2 | + | 40 | 4.68E-11 | 3.69E-08 |
| phosphorus metabolic process (GO:0006793) | 1794 | 22 | 4.32 | + | 5.09 | 4.68E-11 | 3.88E-08 |
| purine ribonucleoside triphosphate biosynthetic process (GO:0009206) | 94 | 8 | 0.23 | + | 35.32 | 1.18E-10 | 8.88E-08 |
| purine nucleoside triphosphate biosynthetic process (GO:0009145) | 95 | 8 | 0.23 | + | 34.95 | 1.28E-10 | 9.18E-08 |
| ribonucleoside triphosphate biosynthetic process (GO:0009201) | 99 | 8 | 0.24 | + | 33.54 | 1.74E-10 | 1.19E-07 |
| nucleoside triphosphate biosynthetic process (GO:0009142) | 110 | 8 | 0.27 | + | 30.18 | 3.84E-10 | 2.52E-07 |
| proton motive force-driven ATP synthesis (GO:0015986) | 67 | 7 | 0.16 | + | 43.36 | 4.89E-10 | 3.08E-07 |
| cellular respiration (GO:0045333) | 176 | 9 | 0.42 | + | 21.22 | 5.46E-10 | 3.31E-07 |
| purine nucleotide biosynthetic process (GO:0006164) | 181 | 9 | 0.44 | + | 20.64 | 6.92E-10 | 4.04E-07 |
| purine-containing compound biosynthetic process (GO:0072522) | 189 | 9 | 0.46 | + | 19.76 | 9.97E-10 | 5.61E-07 |
| ribose phosphate biosynthetic process (GO:0046390) | 193 | 9 | 0.47 | + | 19.35 | 1.19E-09 | 6.46E-07 |
| organophosphate metabolic process (GO:0019637) | 860 | 15 | 2.07 | + | 7.24 | 1.29E-09 | 6.78E-07 |
| aerobic respiration (GO:0009060) | 144 | 8 | 0.35 | + | 23.06 | 2.89E-09 | 1.47E-06 |
| nucleotide biosynthetic process (GO:0009165) | 228 | 9 | 0.55 | + | 16.38 | 4.83E-09 | 2.38E-06 |
| nucleoside phosphate biosynthetic process (GO:1901293) | 234 | 9 | 0.56 | + | 15.96 | 6.01E-09 | 2.87E-06 |
| organonitrogen compound metabolic process (GO:1901564) | 4762 | 31 | 11.47 | + | 2.7 | 7.01E-09 | 3.25E-06 |
| energy derivation by oxidation of organic compounds (GO:0015980) | 245 | 9 | 0.59 | + | 15.25 | 8.83E-09 | 3.98E-06 |
| oxidative phosphorylation (GO:0006119) | 105 | 7 | 0.25 | + | 27.67 | 9.17E-09 | 4.02E-06 |
| purine ribonucleotide biosynthetic process (GO:0009152) | 172 | 8 | 0.41 | + | 19.3 | 1.09E-08 | 4.66E-06 |
| cellular metabolic process (GO:0044237) | 6217 | 35 | 14.98 | + | 2.34 | 1.32E-08 | 5.47E-06 |
| proton motive force-driven mitochondrial ATP synthesis (GO:0042776) | 62 | 6 | 0.15 | + | 40.16 | 1.41E-08 | 5.68E-06 |
| small molecule metabolic process (GO:0044281) | 1574 | 18 | 3.79 | + | 4.75 | 1.46E-08 | 5.74E-06 |
| ribonucleotide biosynthetic process (GO:0009260) | 184 | 8 | 0.44 | + | 18.05 | 1.81E-08 | 6.96E-06 |
| metabolic process (GO:0008152) | 7979 | 38 | 19.22 | + | 1.98 | 1.94E-07 | 7.30E-05 |
| carbohydrate derivative biosynthetic process (GO:1901137) | 538 | 10 | 1.3 | + | 7.71 | 6.18E-07 | 2.27E-04 |
| organonitrogen compound biosynthetic process (GO:1901566) | 1218 | 14 | 2.93 | + | 4.77 | 8.28E-07 | 2.97E-04 |
| organic substance metabolic process (GO:0071704) | 7593 | 36 | 18.29 | + | 1.97 | 1.41E-06 | 4.92E-04 |
| organophosphate biosynthetic process (GO:0090407) | 471 | 9 | 1.13 | + | 7.93 | 1.93E-06 | 6.61E-04 |
| cellular nitrogen compound metabolic process (GO:0034641) | 3235 | 22 | 7.79 | + | 2.82 | 2.18E-06 | 7.30E-04 |
| translational elongation (GO:0006414) | 35 | 4 | 0.08 | + | 47.43 | 2.30E-06 | 7.55E-04 |
| regulation of muscle system process (GO:0090257) | 257 | 7 | 0.62 | + | 11.3 | 3.14E-06 | 1.01E-03 |
| regulation of muscle contraction (GO:0006937) | 166 | 6 | 0.4 | + | 15 | 3.50E-06 | 1.10E-03 |
| cell-substrate junction assembly (GO:0007044) | 40 | 4 | 0.1 | + | 41.5 | 3.76E-06 | 1.16E-03 |
| primary metabolic process (GO:0044238) | 6871 | 33 | 16.56 | + | 1.99 | 4.01E-06 | 1.22E-03 |
| cell-substrate junction organization (GO:0150115) | 41 | 4 | 0.1 | + | 40.49 | 4.12E-06 | 1.23E-03 |
| striated muscle contraction (GO:0006941) | 106 | 5 | 0.26 | + | 19.58 | 6.99E-06 | 2.04E-03 |
| nitrogen compound metabolic process (GO:0006807) | 6312 | 31 | 15.21 | + | 2.04 | 9.73E-06 | 2.79E-03 |
| protein kinase A signaling (GO:0010737) | 15 | 3 | 0.04 | + | 83.01 | 1.04E-05 | 2.94E-03 |
| actin filament-based process (GO:0030029) | 592 | 9 | 1.43 | + | 6.31 | 1.19E-05 | 3.29E-03 |
| cellular nitrogen compound biosynthetic process (GO:0044271) | 1355 | 13 | 3.26 | + | 3.98 | 1.55E-05 | 4.22E-03 |
| cytoskeleton organization (GO:0007010) | 1185 | 12 | 2.86 | + | 4.2 | 2.08E-05 | 5.57E-03 |
| cardiac muscle cell development (GO:0055013) | 71 | 4 | 0.17 | + | 23.38 | 3.19E-05 | 8.37E-03 |
| cardiac cell development (GO:0055006) | 77 | 4 | 0.19 | + | 21.56 | 4.31E-05 | 1.11E-02 |
| actin cytoskeleton organization (GO:0030036) | 538 | 8 | 1.3 | + | 6.17 | 4.46E-05 | 1.14E-02 |
| skeletal muscle contraction (GO:0003009) | 29 | 3 | 0.07 | + | 42.93 | 6.19E-05 | 1.55E-02 |
| phosphorylation (GO:0016310) | 923 | 10 | 2.22 | + | 4.5 | 6.53E-05 | 1.61E-02 |
| respiratory electron transport chain (GO:0022904) | 87 | 4 | 0.21 | + | 19.08 | 6.81E-05 | 1.65E-02 |
| biosynthetic process (GO:0009058) | 2313 | 16 | 5.57 | + | 2.87 | 7.46E-05 | 1.78E-02 |
| muscle cell development (GO:0055001) | 179 | 5 | 0.43 | + | 11.59 | 7.95E-05 | 1.87E-02 |
| organelle organization (GO:0006996) | 2886 | 18 | 6.95 | + | 2.59 | 8.79E-05 | 2.04E-02 |
| regulation of ion transport (GO:0043269) | 769 | 9 | 1.85 | + | 4.86 | 8.91E-05 | 2.03E-02 |
| electron transport chain (GO:0022900) | 98 | 4 | 0.24 | + | 16.94 | 1.06E-04 | 2.39E-02 |
| mitochondrion organization (GO:0007005) | 452 | 7 | 1.09 | + | 6.43 | 1.09E-04 | 2.42E-02 |
| nucleobase-containing compound biosynthetic process (GO:0034654) | 793 | 9 | 1.91 | + | 4.71 | 1.12E-04 | 2.46E-02 |
| monosaccharide catabolic process (GO:0046365) | 36 | 3 | 0.09 | + | 34.59 | 1.13E-04 | 2.43E-02 |
| carbohydrate catabolic process (GO:0016052) | 101 | 4 | 0.24 | + | 16.44 | 1.19E-04 | 2.53E-02 |
| small molecule catabolic process (GO:0044282) | 324 | 6 | 0.78 | + | 7.69 | 1.36E-04 | 2.87E-02 |
| cardiac muscle cell differentiation (GO:0055007) | 106 | 4 | 0.26 | + | 15.66 | 1.42E-04 | 2.95E-02 |
| cardiac muscle tissue development (GO:0048738) | 204 | 5 | 0.49 | + | 10.17 | 1.45E-04 | 2.96E-02 |
| sarcomere organization (GO:0045214) | 41 | 3 | 0.1 | + | 30.37 | 1.62E-04 | 3.27E-02 |
| striated muscle tissue development (GO:0014706) | 210 | 5 | 0.51 | + | 9.88 | 1.65E-04 | 3.25E-02 |
| regulation of system process (GO:0044057) | 651 | 8 | 1.57 | + | 5.1 | 1.65E-04 | 3.29E-02 |
| organic substance biosynthetic process (GO:1901576) | 2240 | 15 | 5.4 | + | 2.78 | 1.93E-04 | 3.75E-02 |
| nucleobase-containing compound metabolic process (GO:0006139) | 2510 | 16 | 6.05 | + | 2.65 | 1.95E-04 | 3.75E-02 |
| actomyosin structure organization (GO:0031032) | 117 | 4 | 0.28 | + | 14.19 | 2.05E-04 | 3.90E-02 |
| heterocycle biosynthetic process (GO:0018130) | 864 | 9 | 2.08 | + | 4.32 | 2.12E-04 | 3.98E-02 |
| musculoskeletal movement (GO:0050881) | 46 | 3 | 0.11 | + | 27.07 | 2.23E-04 | 4.14E-02 |
| multicellular organismal movement (GO:0050879) | 46 | 3 | 0.11 | + | 27.07 | 2.23E-04 | 4.10E-02 |
| positive regulation of transport (GO:0051050) | 1076 | 10 | 2.59 | + | 3.86 | 2.27E-04 | 4.12E-02 |
| aromatic compound biosynthetic process (GO:0019438) | 876 | 9 | 2.11 | + | 4.26 | 2.35E-04 | 4.21E-02 |
| glycolytic process (GO:0006096) | 47 | 3 | 0.11 | + | 26.49 | 2.37E-04 | 4.20E-02 |
| cellular chemical homeostasis (GO:0055082) | 518 | 7 | 1.25 | + | 5.61 | 2.49E-04 | 4.36E-02 |
| muscle structure development (GO:0061061) | 518 | 7 | 1.25 | + | 5.61 | 2.49E-04 | 4.31E-02 |
| ATP generation from ADP (GO:0006757) | 48 | 3 | 0.12 | + | 25.94 | 2.52E-04 | 4.31E-02 |
